# Red blood Cell-derived Extracellular Vesicles as biomaterials: the opportunity of freezing-induced accelerated aging

**DOI:** 10.1101/2025.08.08.669262

**Authors:** Lucia Paolini, Miriam Romano, Valentina Mangolini, Selene Tassoni, Shuhan Jiang, Elena Laura Mazzoldi, Angelo Musicò, Andrea Zendrini, Anna Kashkanova, Vahid Sandoghdar, Anna C. Berardi, Silvia Clara Giliani, Paolo Bergese, Annalisa Radeghieri

## Abstract

Red blood cell-derived extracellular vesicles (RBC-EVs) are emerging as promising biomaterials for next-generation drug delivery, due to their inherent biocompatibility, immune evasion capabilities, and minimal oncogenic risk. However, their clinical translation remains limited by unresolved challenges related to heterogeneity, reproducibility, and long-term storage.

This study proposes a method that leverages freezing-induced ageing for obtaining highly homogeneous RBC-EV batches, an important step towards using RBC-EVs as healthcare biomaterials and advancing their clinical translation in EV-based nanomedicine.

This method was made possible thanks to the analytical support of discontinuous sucrose density gradient and high-resolution interferometric nanoparticle tracking analysis, which allowed the identification of a bimodal subpopulation distribution, in terms of vesicle size, interferometric contrast, and subpopulation profiles, in freshly prepared samples, and then tracked how long-term cold storage at -80 °C channeled this heterogeneity into a monomodal population.

Finally, we evaluated the functionality of homogenized RBC-EV samples by assessing surface-associated enzymatic activity and uptake in cancer cell lines, demonstrating that freeze-thaw-induced accelerated-aging provides a viable strategy for producing RBC-EV preparations that retain membrane integrity and remain readily internalized by cells.

These findings offer valuable insights into the optimization and standardization of RBC-EV handling and storage protocols, providing a foundation for their reliable integration into EV-based therapeutic applications.

## 1. Introduction

Extracellular vesicles (EVs) are a groundbreaking biological discovery of the past two decades. These natural lipid nanoparticles, ranging in size from 30 nm to 1000 nm, are secreted by all cell types in animals and bacteria, acting as carriers for many biomolecules^1^. EVs play a vital role in mediating physiological processes and contributing to disease spread, including cancer^2^ and infections^3^. Their unique properties, which have evolved for efficient transport and pharmacokinetics, make them highly promising for precision nanomedicine^4^. Early clinical trials of EV-based therapeutics have shown encouraging results, fueling ongoing research into their potential as novel biomaterials i.e. as drug delivery vehicles^5,6^.

Red Blood Cell-derived EVs (RBC-EVs) stand out as particularly promising biomaterials for next-generation drug delivery^6^ RBC-EVs offer several key advantages: they are devoid of mitochondrial and nuclear DNA, thus posing minimal oncogenic risk, and exhibit excellent biocompatibility^7–10^. RBC-EVs can be produced at a large-scale with good reproducibility and benefit from well-established protocols in transfusion medicine^11^. Additionally, RBC-EVs derived from patients’ blood offer precision and safety, making them an attractive option for clinical translation in drug delivery and therapeutic applications.

RBC-EVs large-scale *in vitro* production can be boosted by exploiting external stimuli (i.e., physical or chemical stress)^12^. The main chemical inductors used so far in this context are calcium ionophores, as Ionomycin and A23187^13^, lysophosphatidic acid (LPA)^14^, and phorbol-12 myristate-13 acetate (PMA)^15^. This procedure is then followed by the separation of RBC-EVs from RBCs by different methods, which usually include serial ultra-centrifugation steps and a sucrose gradient^8^.

Compared to naturally secreted RBC-EVs obtained after RBC storage^16,17^ up to 28 days at +4°C, chemically induced RBC-EVs show a very similar enrichment of EV markers with size and shape typical of EVs^8,18^. Furthermore, the biochemical composition obtained by proteomic analysis looks very similar apart from the enrichment of calcium-binding proteins^19^ and some differences in membrane protein expression^20^.

Despite earlier findings^18^, it has recently been shown that fresh chemically induced RBC-EVs exhibit two separate size and density profiles, each with distinct protein expression patterns. These variations lead to differing physiological responses upon internalization^21^. In addition, it has been demonstrated that different storage conditions can impact EV yield, physical properties, and functionality^22^, and combined technologies are needed to characterize all the modifications occurring in EV samples^23^.

Thus, in order to enable biobanking of RBC-EVs for large-scale production and use, it is essential to conduct extensive studies on their homogeneity and stability after different storage conditions. These factors are crucial as different storage protocols can significantly influence the biological activity of RBC-EVs^24–26^. Additionally, to ensure the therapeutic applicability of RBC-EVs in large quantities, it is necessary to evaluate whether their heterogeneity persists during storage and whether maintaining this heterogeneity is beneficial. For clinical use, however, a more homogeneous product may be preferable to ensure consistent therapeutic outcomes^27^ according to international medicines regulatory agencies guidelines^28,29^.

In order to achieve this, key technologies at the single-vesicle or subpopulation-level analysis have been developed as high-sensitivity flow cytometry^30^, NanoFlow Cytometry (NanoFCM)^31^, and which enable size and surface marker profiling. ExoView^32^, a chip-based immunocapture platform, allows multiplexed phenotyping of individual EVs without extensive preprocessing. Structural heterogeneity can be explored via atomic force microscopy (AFM)^33^ [] and electron microscopy (EM)^34^ [], which provide nanoscale resolution. Raman and FT-IR spectroscopy^35,36^ [] offer label-free biochemical fingerprints, while surface plasmon resonance (SPR)^37^ [] detects molecular interactions in heterogeneous EV populations.

Recently, interferometric nanoparticle tracking analysis (iNTA) has been introduced in this scenario with the advantage of high-precision measurement of EV size and content by refractive index determination^38,39^. Employing at least two orthogonal analytical methods is widely recommended to minimize methodological bias.

In this study, by combining a multiparametric characterization based on iNTA and discontinuous density gradient (DSG), we developed an “accelerated-aging” protocol based on freeze-thaw cycles that allows to obtain highly homogeneous RBC-EV preparations. These preparations were assessed for RBC-EV membrane preservation through RBC-EV surface associated enzymatic assay and for their capacity for RBC-EVs to be uptaken from cells.

## 2. Materials and Methods

We have submitted all relevant data from our experiments to the EV-TRACK knowledge base (EV-TRACK ID: EV250035)^40^.

### RBC-EV production

RBC-EVs were isolated according to the protocol outlined by previous works^8,18^. In brief, upon the collection of 100 ml of blood, red blood cells (RBCs) were separated by centrifugation at 1000 × g for 8 min at 4 °C and underwent three washes in PBS without calcium and magnesium. Following this, RBCs were subjected to two additional washes with CPBS (PBS + 0.1 g/L calcium chloride) and then transferred into a 75 mm^2^ tissue culture flask. Calcium ionophore was added to the flask at a final concentration of 10 µM, and the mixture was incubated overnight at 37 °C. Subsequently, 75 ml of RBCs were gently collected from the flask, and any cellular debris was eliminated through a series of differential centrifugation steps (600 × g for 20 min, 1600 × g for 15 min, 3260 × g for 15 min, and 10,000 × g for 30 min at 4 °C). At each step, the pellet was discarded, and the supernatant was transferred into a fresh tube. The resulting supernatants were filtered through a sterile 0.45 μm nylon syringe filter. EVs were obtained through ultracentrifugation at 50,000 × g for 70 min at 4°C. The EV pellets were then resuspended in cold PBS, layered above a 2 ml frozen 60% sucrose cushion, and centrifuged at 50,000 × g for 16 h at 4 °C, with the deceleration speed set to 0. The red layer of EVs was collected, washed twice with cold PBS, and spun down at 50,000 × g for 70 min at 4 °C. Finally, the EVs were resuspended in 1 ml of cold PBS and stored at 4°C (Fresh samples), or at -80°C for 6 months (6-month samples), or at -80°C for 1 year (1-year samples) for the further analyses. Red Blood Cells were obtained from five anonymized healthy volunteers under written consent and provided by the A. O. Spedali Civili di Brescia, ethical approval “EritrEV NP5705”.

### Freeze-and-thaw cycles

The Fresh RBC-EV preparations, stored at 4°C (maximum time 1 week), were subjected to two types of freeze-and-thaw cycles: Light_5 cycles at -80°C and 37°C (water bath), 5 min each incubation; Strong_10 cycles at -80°C and 37°C (water bath), 5 min each incubation, before further analyses.

### Bicinchoninic acid (BCA) and Bradford assay

Protein concentrations of samples were determined with Pierce™ BCA Protein Assay Kit (ThermoFisher, Rockford, USA) and Bradford assay kit (Biorad) following the manufacturer’s instructions.

### Discontinuous Density Gradient (DSG)

DSG was carried out by adapting the protocols developed in previous studies^41,42^. RBC-EVs produced as previously described were quantified using the Bradford assay, then 600 ug were centrifuged at 50,000 x g at 4°C for 70 min. Briefly, RBC-EV preparations were resuspended in buffer A (10 mM Tris–HCl, 250 mM sucrose, pH 7.4) to a final volume of 1 ml, loaded on top of a discontinuous sucrose density gradient (15% (600 μl), 20%, 25%, 30%, 40%, 60%, 65% (400 μl), 70% (800 μl) sucrose in 10 mM Tris–HCl, pH 7.4) and centrifuged at 230,000 x g for 16 h at 4°C (rotor MLS 50; Beckman Optima MAX, no brake). Twelve fractions with equal volumes (400 μl) were collected from the top of the gradient. All fractions were diluted with 600 μl water suitable for high-performance liquid chromatography (HPLC water, final volume 1 ml) and ultracentrifuged at 100,000 x g for 2 h at 4°C (Optima MAX, TLA–55 rotor, 1.5 ml polypropylene microfuge tube, Beckman). Pellets were resuspended in 100 μl HPLC water previously ultracentrifuged at 100,000 g for 2 h in Optima MAX, TLA–55 rotor, 1.5 ml polypropylene microfuge tubes (Beckman) and aliquoted for further analyses.

### SDS-PAGE and Western Blot analysis

SDS−PAGE and Western blot were performed by standard procedures^43,44^. Both RBC-EV homogenate (H) and RBC-EV gradient fractions were denaturated with a 6X Laemmli buffer and boiled for 5 min at 95°. For RBC-EV homogenate, 30 µg were loaded, while for gradient fractions 15 μl each fraction were loaded. Samples were electrophoresed on acrylamide-bisacrylamide gels 12.5% and analyzed by Western Blot with the following antibodies (at dilution 1:500): mouse anti-Flotillin 1 (Santa Cruz, clone C-2, sc-74566), rabbit anti-ADAM10 (Origene, AP05830PU-N), mouse anti CD81 (Santa Cruz Biotechnology, clone B11, sc-166029), mouse anti CD9 (Santa Cruz Biotech, clone C-4, sc-13118), mouse anti-TSG101 (Santa Cruz Biotechnology, clone C–2, sc–7964), mouse anti LAMP1 (BD Biosciences, 611042/611043), mouse anti-Annexin V (Santa Cruz Biotech, clone H-3, sc-74438), rabbit anti-AnnexinXI (Genetex, CTX33010), mouse anti Alix (Santa Cruz Biotech, clone 2H12, sc-53539), mouse anti CD45 (Santa Cruz Biotechnology, clone 35-Z6, sc-1178), mouse anti Integrin αIIb/ITGA2B/CD41 (Santa Cruz, clone B-9), mouse anti-GAPDH (Millipore, clone 6C5), mouse anti-Band 3 (Santa Cruz Biotech, clone A-6, sc-133190). Mouse anti-Hemoglobin A (Abnova, clone 4F9) and Mouse anti-Hemoglobin B (Abnova, colone 7B12) were diluted 1:1000. The blots were detected using a Luminata Classic HRP Western substrate (Millipore). The images were acquired using a G:Box Chemi XT Imaging system, as previously described^45,46^.

### UV-VIS spectroscopy

RBC-EV gradient fractions were collected and treated as described above. Resuspended pellets were analyzed for the presence of hemoglobin by measuring the absorbance at 414 nm with UV-VIS spectroscopy Nanodrop spectrophotometer (Thermo Scientific) as previously described^47^.

### Colorimetric nanoplasmonic (CONAN) assay

The purity of RBC-EV preparations from soluble contaminants was tested with the CONAN assay according to Zendrini et al^48,49^. The assay is a colorimetric test that exploits the aggregation of citrate-capped gold nanoparticles (AuNPs) onto the EV membrane and the formation of the protein corona on the AuNP surface to detect soluble proteins in EV preparations^49^. The result of the assay was collected on Ensight Multi Mode Reader (Perkin Elmer). Measurements were performed for each sample in triplicate and expressed as the mean of three replicates ± standard deviation.

### Atomic Force Microscopy (AFM)

Atomic force microscopy (AFM) imaging was performed with a Nanosurf NaioAFM equipped with a Multi75-AI-G tip (Budget Sensors). For sample preparation, RBC-EVs (bulk) pellets were resuspended in 100 μL sterile H_2_O (Milli-Q, Merck Millipore) and diluted 1:10 in H_2_O. Five μL of the sample were then spotted onto freshly cleaved mica sheets (Grade V-1, thickness 0.15 mm, size 15 × 15 mm^2^) and air-dried at 37 °C for 10 min. Images were acquired in tapping mode, with a scan size ranging from 1.5 to 25 mm and a scan speed of 1 s per scanning line^36^. Image processing was performed on Gwyddion ver. 2.61.

### Nanotracking particle analysis (NTA)

RBC-EV preparations were resuspended in 100 μl of sterile HPLC water (Milli-Q; Merck Millipore) for the characterization by Nanoparticles Tracking Analysis (NTA) using a NanoSight NS300 system (Malvern, Panalytical LTD, Malvern, UK) to evaluate the concentration and size distribution of EVs. The system was equipped with a Blue488 laser and sCMOS camera. Before sample analysis, filtered PBS (0,22μm filtered) was analysed for particle contamination. For the analysis, all samples were diluted in filtered PBS (1:1000) to a final volume of 1 ml to obtain optimal particle number per frame value (20-120 particles/frame). EVs were injected in the sample chamber through a Nanosight syringe pump (Malvern, Panalytical LTD, Malvern, UK) that provides a continuous flow (50μl/min) using a 1 ml syringe at room temperature. Recordings of the movements of particles were collected for 60s, five times for each sample at a camera level of 12 using the best detection threshold. The mean, mode, and median EV sizes from each video were used to calculate the sample concentration, expressed in particles per ml^50^.

### Interferometric Nanoparticle Tracking analysis (iNTA)

Samples (at least 3 biological replicates for each point analyzed) were diluted 1:100 and analyzed using the iNTA instrument^38^. An iNTA measurement setup was utilized for particle analysis^51^. Measurements were conducted using IBIDI μ-Slide 18 well-chambered cover glasses, which were plasma cleaned and passivated with an mPEG2000-Silane (Laysan Bio) solution^39^. The microscope focus was adjusted to 1 μm above the cover glass, and 100 μL of diluted RBC-EVs were measured over 30 min. Measurements were performed at 10 kHz with a 50 μs exposure time. The field of view (FOV) is 6 × 6 μm2. Particle size was derived from the diffusion constant. Contrast^1/3^ was calculated as a ratio of scattred field to reflected field.

### MDA-MB-231 cell culture

MDA-MB-231 cell line was purchased from the American Type Culture Collection (ATCC; Manassas, VA, USA). For culture maintenance, cells were cultured in Dulbecco’s Modification of Eagle’s Medium (DMEM) with 4.5 g/L glucose, L-glutamine & sodium pyruvate (Corning, USA) supplemented with 1% penicillin-streptomycin and 10% Fetal Bovine Serum (FBS, EuroClone S.p.a, Italy).

### RBC-EV labelling

RBC-EV labeling with MemGlow™ 488 was performed following standard customer protocols as previously described^8^. Briefly, RBC-EV Fresh, 6-month, Strong F-T samples (10^11^ particles each sample) were incubated with 5 μl MemGlow™ 488 100 nM for 15 min at RT (final volume 1 ml in PBS), and then excess MemGlow™ 488 was removed by ultracentrifugation (100.000×g, 2 h, Optima max -XP equipped with a TLA-55 rotor, Beckman Coulter, USA). Before cell treatments, pellets were resuspended in 500 μl DMEM serum free medium.

### Flow cytometry

MDA-MB-231 cells were seeded on a 12-well plate, 4×10^5^ cells/well, in 1 ml of complete media. After 24 h, MDA-MB-231 cells were washed twice with PBS and incubated for either 30 min or 4h with MemGlow™ 488-stained RBC-EVs-Fresh/6months/F-T strong (10^11^ RBC-EV per well, 500 μl final volume), unstained RBC-EVs Fresh (10^11^ RBC-EV), or with MemGlow™ 488 fluorescent dye without RBC-EVs. The treatment with MemGlow™ 488 fluorescent dye without EVs was done to exclude any possible unspecific fluorescent signal. Untreated cells were used as a negative control. After incubation, cells were washed with PBS and treated with 200 μl 0.25%trypsin-EDTA (Corning, USA) for 1 min at 37°C, both to remove externally bound EVs and to detach cells from the culture vessel. After trypsin treatment cells were resuspended in 500 μl of DMEM medium supplemented with 5% Fetal Bovine Serum and centrifuged at 800xg for 5 min. Pellets were resuspended in 200 μl DMEM Serum Free and analyzed by flow cytometry. Flow Cytometry acquisition was performed on FACSCanto™ II (BD, Franklin Lakes, NJ) from at least 2×10^4^ events/tube, excluding debris/dead cells and doublets. Data were elaborated with FlowJo v10.6.2 software (TreeStar, Ashland, OR). The results were expressed as the difference between the median fluorescence intensity of cells incubated with MemGlow™ 488-stained RBC-EVs and of the corresponding untreated control (ΔMFI).

### Acetylcholinesterase activity assay

Acetylcholinesterase activity was assessed by standard procedures^52,53^. Briefly, 3×10^9^ particles of RBC-EVs Fresh, 6-month at -80°C and after Strong F-T treatment, were diluted in 60 μl of PBS without calcium and magnesium and incubated with 1.25 mM of acetylthiocholine (Sigma-Aldrich) and 0.1 mM of 5,5’-dithio-bis(2-nitrobenzoic acid) in a final volume of 1 ml. The incubation was carried out in 96-well plates at 37°C, and the change in absorbance at 415 nm was followed at 0, 10, 20, 30, and 60 min. Absorbance was measured with the spectrophotometer model 680 Bio-Rad, USA). Data are presented as the mean of three technical replicates ± Standard Deviation (SD).

### Statistical Analysis

For the iNTA and flow cytometry analyses, statistical analysis was made using Student t-test: ** p value ≤ 0.01, *** p value ≤ 0.001, **** p value ˂ 0.0001, ns = not significant. For the measurements of the mean diameters of the RBC-EV fractions calculated with the NTA, the statistical analysis was performed with ordinary one-way ANOVA. ****p value < 0.0001. All statistical analyses were performed using Graph Pad Prism software.

## 3. Results and Discussion

### 3.1 Fresh RBC-EV production and characterization

Red Blood Cell-derived EVs (mean concentration 9 x 10^11^± 3 x 10^9^ RBC-EV/ml measured by NTA) were obtained following the protocol described by Usman et al^18^. from the RBCs of five healthy volunteers and stored at +4°C for maximum 1 week (Fresh). Preparations were characterized according to the most updated guidelines^54^. In particular, Western blot analysis was used to identify the presence or absence of specific protein markers. Specifically, RBC-EVs were found to express Band 3, GAPDH, Hemoglobin subunit β (HBB), Hemoglobin subunit α (HBA), Flotillin-1, Annexin-V, LAMP-1, Annexin-XI, and ADAM10. Among these, Band 3, HBB, and HBA are specific markers of RBC-EVs, while the others represent common markers of EVs^1,11,54^. In contrast, the RBC-EV preparations did not express CD45, a typical marker of leucocytes, nor CD41, a platelet-specific marker, indicating that the RBC-EV preparations were free from contamination by other blood components^55^. Additionally, the RBC-EV lacked the tetraspanins CD9 and CD81, which aligns with findings in the existing literature^56^ (Fig.1A). Bradford assay and Colorimetric Nanoplasmonic (CONAN) Assay^48,49^ (Fig. Supplementary S1A-B) were used to determine protein concentration and the absence of soluble protein contaminants, respectively, while Atomic Force Microscopy (AFM) was implemented to investigate the morphology of the nano-objects in the preparations (Supplementary Fig. S1C). Data were in accordance with our previous works^8,57^. In addition, the samples were analyzed for their nanoparticle size distribution using Nanoparticle Tracking Analysis (NTA). Results shown in Fig. 1B highlighted that, with this analytical technique, we cannot appreciate significant differences in terms of subpopulation heterogeneity since all samples showed a median homogeneous distribution around 165 ± 13.2 nm. These data were in contrast with previous findings about freshly isolated RBC-EVs kept at +4°C^21^, indicating that, probably, conventional NTA alone may not be the best option to investigate subpopulation heterogeneity. Therefore different methods with higher sensitivity and better resolution are needed^38,58^.

**Fig. 1.**
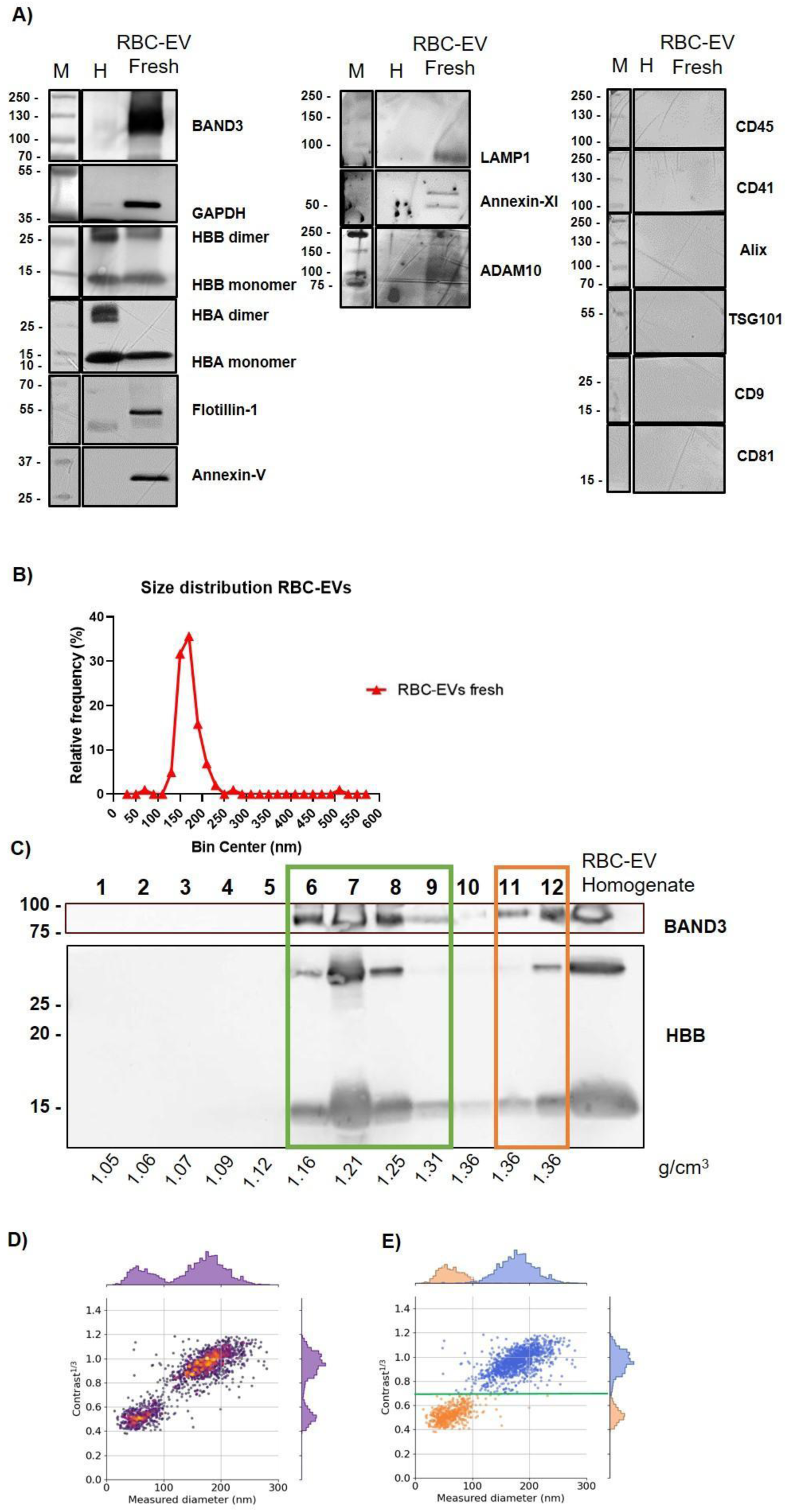
RBC-EV Fresh characterization. A) Biochemical characterization of RBC-EV RBC-EVs stored at 4°C for 1 week (Fresh) preparations; M = marker, H = cellular homogenate 30µg. B) Size distributions of RBC-EVs Fresh. Size distributions obtained by Nanoparticle Tracking Analysis (NTA). C) Identification of two subpopulations with Western Blot of Discontinuous Sucrose Gradient (DSG)-fractions of RBC-EVs Fresh preparations. Protein detected are Band 3 anion transport protein (BAND3) and Hemoglobin subunit β. For RBC-EV homogenate loaded 30 µg. Green box: RBC-EV fraction with density from 1.16 to 1.31 g/cm3; orange box: RBC-EV fraction with density of 1.36 g/cm3. D) Size distribution and contract measurements of RBC-EV Fresh obtained with iNTA E) Identification of two RBC-EV Fresh subpopulations with different size and contrast by iNTA. Orange dots EVs with size from 30 to 100 nm and contrast below 0.7; Blue dots EVs with size from 101 to 300 nm and contrast from 0.71 to 1.4.

### 3.2 Orthogonal methods reveal heterogeneity in freshly isolated RBC-EVs

In order to investigate heterogeneity of RBC-EV preparations two different bio-orthogonal techniques were introduced: discontinuous sucrose density gradient (DSG) and interferometric NTA (iNTA). According to recent literature, different RBC-EV populations can be differentiated with more precise optical methods^21^. We, thus, decided to perform iNTA analysis on the RBC-EV preparations. This method allowed us to differentiate nanoparticles by their size and interferometric contrast (contrast ^1/3^). In addition, this method has been demonstrated to have a higher sensitivity and better resolution to determine the size distribution of small nanoparticles compared to conventional NTA^38,39^.

Fresh RBC-EV preparations were loaded on top of a DSG separately and centrifuged for 16h at 230.000 x g. Twelve fractions of equal volume (400 μl) were collected, processed as described in Materials and Methods, and analyzed for their biomolecular profile, density^41,59^, particle number and size distribution. As shown in Fig. 1C, Western Blot analysis for typical RBC-EV markers Band 3 and HBB^18,21,33^ of the 12 fractions revealed the presence of two subpopulations: one distributed within fractions bearing density from 1.16 to 1.31 g/cm^3^ (fractions 6-9, green box) and the other with density around 1.36 g/cm^3^ (fractions 11 and 12, orange box). Band 3 and HBB signal, both in the monomeric form (detected at 15 kDa) and the dimeric form (detected at 30 kDa), were perfectly matching among the fractions. The hemoglobin distribution of RBC-EV in DSG fractions was also analyzed by UV-VIS spectroscopy absorbance at 414 nm. Results confirmed the distribution highlighted in the Western Blot with a peak of absorption in fractions 7 (Supplementary Fig.S2A). NTA analysis of DSG fractions confirmed an enrichment of nanoparticles in fractions 6-7 (Supplementary Fig. S2B), and the BCA assay confirmed the presence of proteins, especially in fraction 7 (Supplementary Fig. S2C).

Measurements on Fresh samples obtained with iNTA aligned perfectly with the DSG distribution, even though the two methods assess different physical and chemical properties since DSG focuses on density, while iNTA measures contrast and size distribution.

As shown in Fig 1D and 1E, RBC-EVs showed a bi-modal distribution when contrast ^1/3^ is plotted *vs.* hydrodynamic size for each nanoparticle. We can demonstrate the presence of two subpopulations with contrast^1/3^ below 0.7 and size distribution mostly below 100 nm (Fig. 1E, orange dots, Lower subpopulation) and another with contrast^1/3^ above 0.7 and size distribution mostly above 100 nm (Figure 1E, blue dots, Upper subpopulation).

### 3.3 Long-term storage conditions at -80°C affect RBC-EV heterogeneity

With the objective to homogenize RBC-EVs towards clinical translation we checked if short or long term storage at -80C could have an effect on this parameter. We conserved RBC-EV preparations for 6 months at -80°C (6-month) or 1 year at -80°C (1-year).

We next evaluated subpopulation heterogeneity, applying the same orthogonal methods as for the Fresh samples. First, compared to Fresh sample (Fig 2A), the 6-month sample showed a different distribution among the DSG fractions in Western Blot: protein signals in fractions at higher density (11 and 12) were no longer present (Fig.2B). Instead, both Band 3 and HBB remained distributed in fractions 6-8, with an additional presence in lighter fraction, particularly fraction 4 and 5 (density 1.09-1.12 g/cm^3^). This shift was especially noticeable for the Band 3 signal (detected at 95 kDa) and the monomeric form of hemoglobin (detected at 15 kDa) (Fig. 2B Blue box). These results were confirmed through spectroscopic analysis of the hemoglobin content that showed a peak of absorption in fraction 5. (Supplementary Fig. S2A). NTA measurements indicated an enrichment of nanoparticles in fractions 4 and 5 (Supplementary Fig.S2B), at higher concentrations with respect to the Fresh samples. Likewise, the BCA assay confirmed a higher protein concentration in fraction 5 respect to Fresh sample (Supplementary Fig. S2C).

**Fig. 2.**
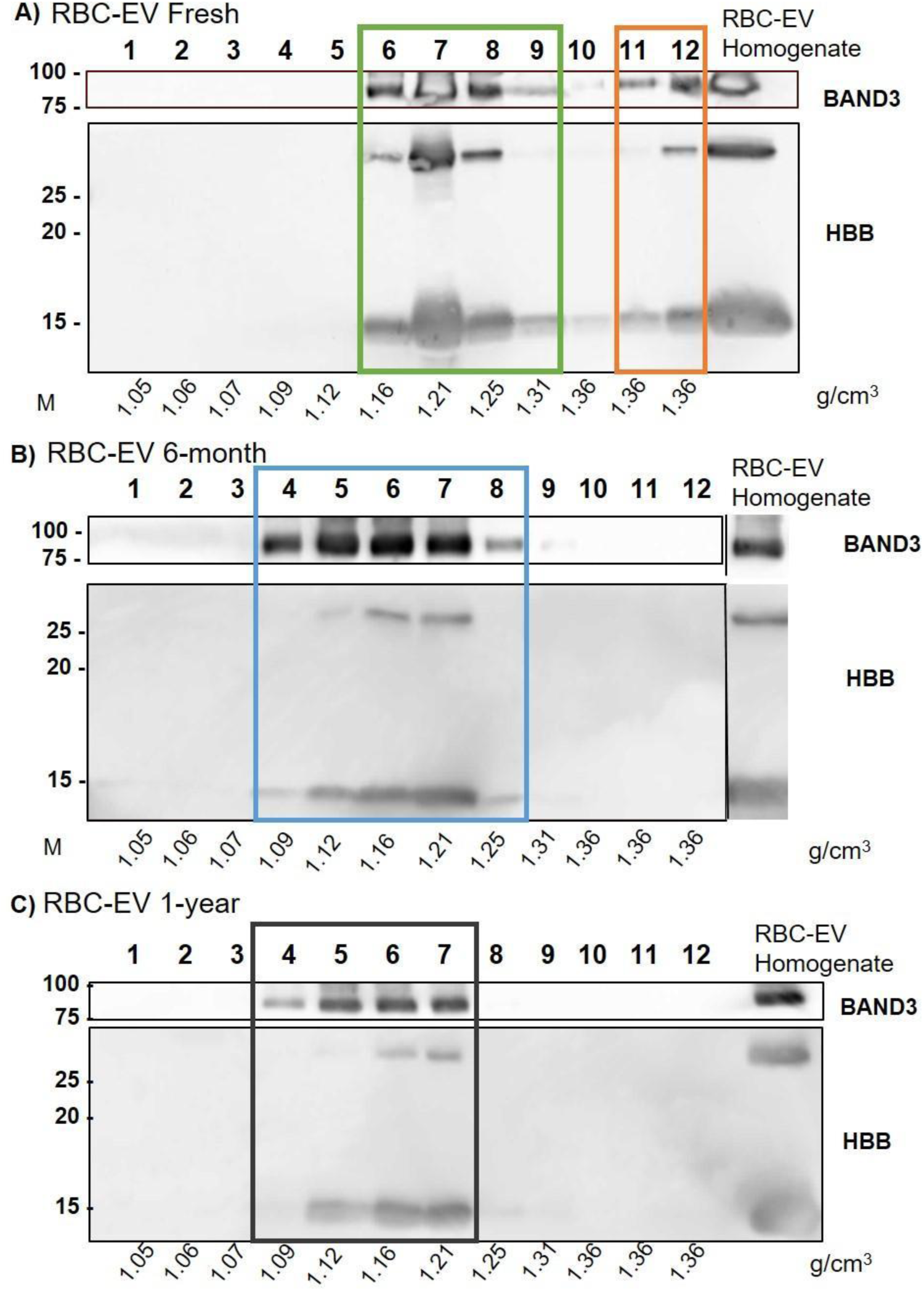
Gradient distribution of RBC-EVs after different storage conditions. **A)** Western Blot of RBC-EVs after DSG; equal volume (15 μL) of each fraction (1-12) for Fresh **(B)**, 6-months **(C)**, 1-year were loaded; 30 μg of RBC-EV homogenate samples were loaded. Samples were electrophoresed on SDS-PAGE gel (12% (Acrylamide/Bis-acrylamide) and analyzed for the antibodies described in the figures. Un-cropped WB are available in the Supplementary Information (Supplementary Figure S3). Each fraction density is indicated in g/cm^3^. RBC-EV subpopulations with different gradient distributions are indicated in colored boxes: A) green box fractions 6-9; orange fractions 11-12; B) blue box fractions 4-9; C) black box 4-7.

A similar trend was observed for the 1-year sample, which showed an even more pronounced narrowing of the RBC-EV distribution in the DSG fractions (from fractions 4 to 7; corresponding to a density of 1.09-1.21 g/cm^3^) (Fig. 2C, black box) and a consequential disappearance of the Band 3 signal from fractions 8 and 9, and a reduction of the hemoglobin signal in these two fractions. Uncropped version of the Western blot images described so far can be found in Supplementary Figure S3A,B,C.

Spectroscopic analysis of the hemoglobin content aligned perfectly with these observations, confirming a decrease of the hemoglobin in these two fractions (Supplementary Fig. S2A) and an increase signal in fraction 5 and 6 respect to Fresh. NTA and protein measurements confirmed the results with an increased concentration of nanoparticles in fractions 4 and 5, in fractions 5 and 6 respectively, compared to Fresh sample (Supplementary Fig. S2B-C).

In addition, NTA analysis of the DSG fractions from 4 to 7 highlighted changes in the size distributions between the three RBC-EV preparations. Indeed, DSG fractions of the Fresh RBC-EV samples showed the presence of two statistically significant subpopulations in fractions 4-5 compared to fractions 6-7 (Supplementary Fig. S4A). Particles in fractions 4-5 present a mean size of 123 ± 51 nm while particles in fractions 6-7 a mean diameter of 170 ± 10 nm (Supplementary Fig. S4B). On the other hand, the 6-month and 1-year samples did not show the dual size distribution in fractions 4-5 and 6-7 observed in the Fresh samples (Supplementary Fig. S4C,D,E,F).

Taken together, these data indicate that the RBC-EV subpopulations undergo significant changes in their distribution and composition during long-term storage, with a shift toward lighter fractions and a reduction in the density and modulation in size of the particles, suggesting potential alterations in RBC-EV heterogeneity over time.

iNTA measurements further corroborated this data, showing that the separation in two distinct subpopulations becomes significantly less prominent for samples that have been stored at -80°C. Nevertheless, we can quantify the relative amount of nanoparticles with contrast^1/3^ above and below 0.7 in all samples.

The Fresh sample (Fig 3A) showed that around 70% of the relative concentration of RBC-EVs were found in the upper population (Fig. 3G, contrast^1/3^ > 0.7, blue dots), while 30% were distributed in the lower population (Fig. 3G, contrast^1/3^ < 0.7, orange dots).

**Fig. 3.**
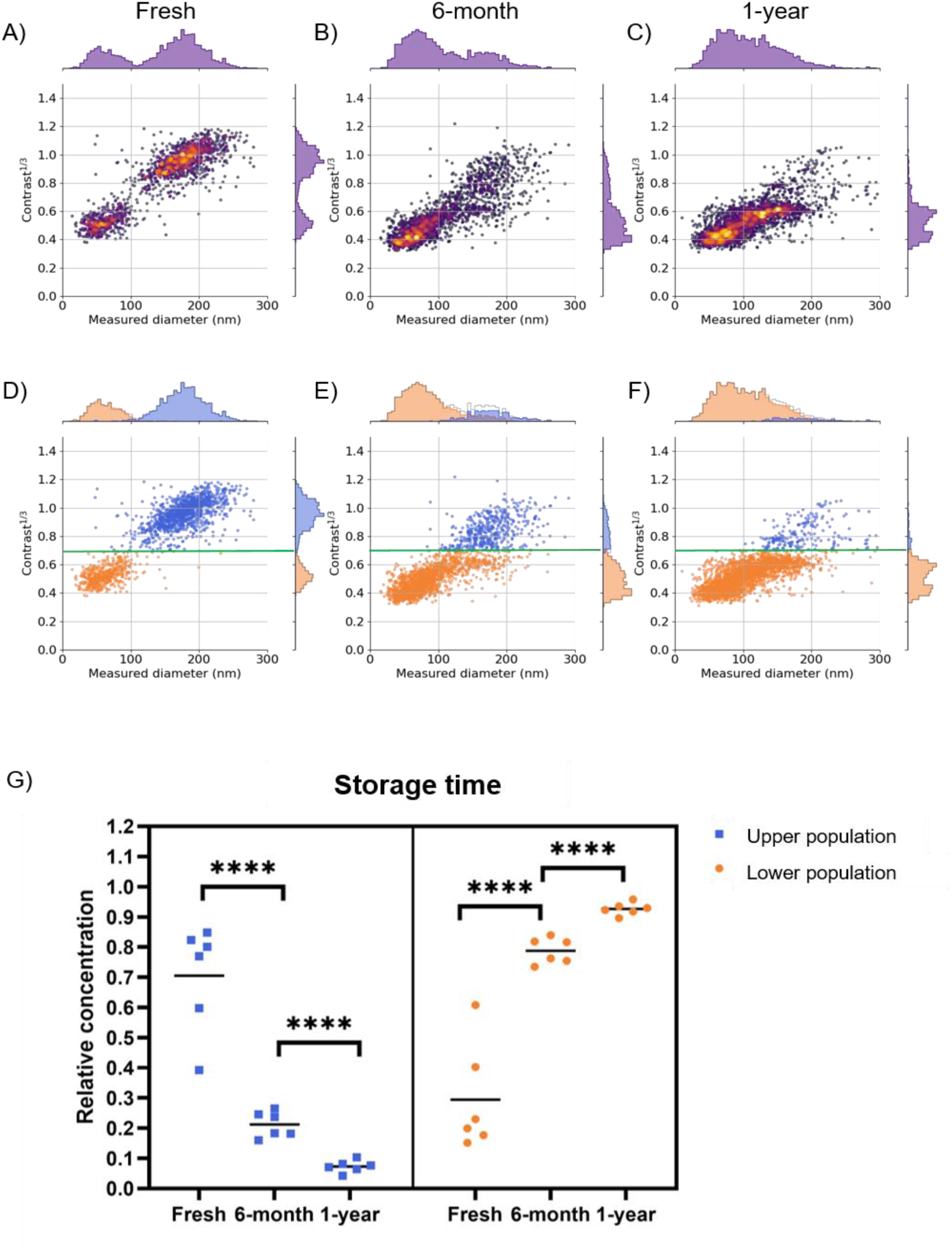
Interferometric NTA (iNTA) analysis. Fresh **(A)**, 6-month **(B),** and 1-year **(C)** samples were analyzed by iNTA. Images represent size distribution (x-axis) and contrast (y-axis) of the nanoparticles. Nanoparticles subpopulations grouped for their contrast in Fresh **(D)**, 6-month **(E)**, and 1-year **(F)** samples. The green line represents a contrast^1/3^ value of 0.7. Nanoparticle distribution determined for contrast^1/3^ ˃ 0.7 (blue dots, Upper population) or ˂ 0.7 (orange dots, Lower population). **G)** Relative Quantification of nanoparticles present in the Upper (contrast ^1/3^ ˃ 0.7) and Lower (contrast ^1/3^ ˂ 0.7) population for Fresh, 6-month, and 1-year samples. Statistical analysis using Student t-test: **** p value ˂ 0.0001.

The 6-month sample showed an opposite distribution compared to the Fresh sample (Fig. 3B, 3E): fewer nanoparticles were distributed in the Upper population (Fig. 3G, contrast^1/3^ > 0.7, blue dots) compared to the Fresh sample, with their relative concentration decreasing to 20% (Fig. 3G, blue dots). Meanwhile, the relative concentration of the lower population (contrast^1/3^ < 0.7) increased to 80% of the total nanoparticles analyzed (Fig. 3G, **** p < 0.0001). This trend was even more pronounced in the 1-year sample: only 10% of nanoparticles were distributed in the upper population, while 90% were found in the lower population. The shift in distribution between the 6-month and 1-year samples was also statistically significant (Fig. 3G, **** p < 0.0001).

These data are in accordance with DSG distribution and confirm that storage reduces the heterogeneity of RBC-EVs. Over time, the two distinct subpopulations gradually merge into a more uniform population in terms of density and contrast^1/3^, with this shift becoming more pronounced in a time-dependent manner.

### 3.3 Freeze-and-thaw cycles decrease RBC-EV heterogeneity

To develop a method that ensures consistent conditions in RBC-EV preparations similar to those found in long-term stored samples, without the need for prolonged storage, we exposed Fresh samples to two different types of freeze/thaw (F-T) cycles. The first treatment, labeled “Light F-T,” involved 5 cycles at -80°C and 37°C, each lasting 5 min. The second treatment, labeled “Strong F-T,” involved 10 cycles at the same temperatures and duration. The samples were then analyzed by DSG to biochemically confirm the changes in the density distributions and with iNTA to assess changes in the relative concentrations in terms of contrast^1/3^ of the subpopulations of RBC-EVs. It is to note that the Light F-T and Strong F-T treatments did not alter significantly the overall nanoparticle concentrations of samples as checked by iNTA (Supplementary Fig. S5A). DSG processing of Light F-T and Strong F-T samples revealed that the two treatments induced a change in the distribution within the gradient fractions in comparison to the Fresh sample. After Light F-T treatment, we could still observe the presence of two subpopulations (Fig. 4A): one from fractions 5 to 9 (blue box, density from 1.11 to 1.31 g/cm^3^) and another in fractions 11 and 12 (orange box, density 1.35 g/cm^3^). Light F-T samples presented a wider distribution in the lighter fractions compared to Fresh samples. Notably, fraction 5 (1.11 g/cm^3^) in Light F-T samples contained both RBC-EV markers, Band 3 and HBB, whereas Fresh samples never reached this density. In the Fresh sample, the lighter fraction containing both Band 3 and HBB was fraction 6, with a density of 1.16 g/cm^3^ (Fig. 1C). Additionally, the HBB present in fraction 5 of the Light F-T sample was also different from the Fresh sample in fraction 6. Fresh sample fraction 6 contained both the monomeric and dimeric form of HBB, according to the molecular weights of the Bands, while Light F-T fraction 5 contained only the monomeric form of HBB. This trend was also visible in lighter fractions of 6-month and 1-year samples (Fig 2B,C). It has been demonstrated that the environment can influence hemoglobin structural changes^60^. We hypothesize that, in our experimental conditions, the monomeric form of HBB could be associated with EVs in a more stable manner than the dimers. Further studies are needed to assess the topology of HBB associated with RBC-EV: if it could be considered a cargo or part of RBC-EV “biomolecular corona^57^”. The trend described for the Light F-T sample was even more pronounced for the Strong F-T sample. We could observe a signal of RBC-EV also in fraction 4 (density 1.09 g/cm^3^) (Fig. 4B, blue box), and both fractions 4 and 5 contained only the monomeric form of HBB. On the other hand, the distribution in the denser fractions also shifted compared to the Light F-T and Fresh samples. Fraction 11 (1.35 g/cm^3^) was no longer populated, and both Band 3 and HBB (monomeric form) were retained only in fraction 12 (Fig. 4B, orange box). Uncropped version of the Western blot images described of Light F-T and Strong F-T can be found in Supplementary Figure S6A,B. It is to note that the DSG factions 4 and 7 of the Strong F-T sample contained intact vesicles as checked by AFM, indicating that the F-T processes did not alter the morphology of nanoparticles and that changes in DSG fractions are not due to enrichment of cellular debris (Supplementary Fig. S5B,C).

**Fig. 4.**
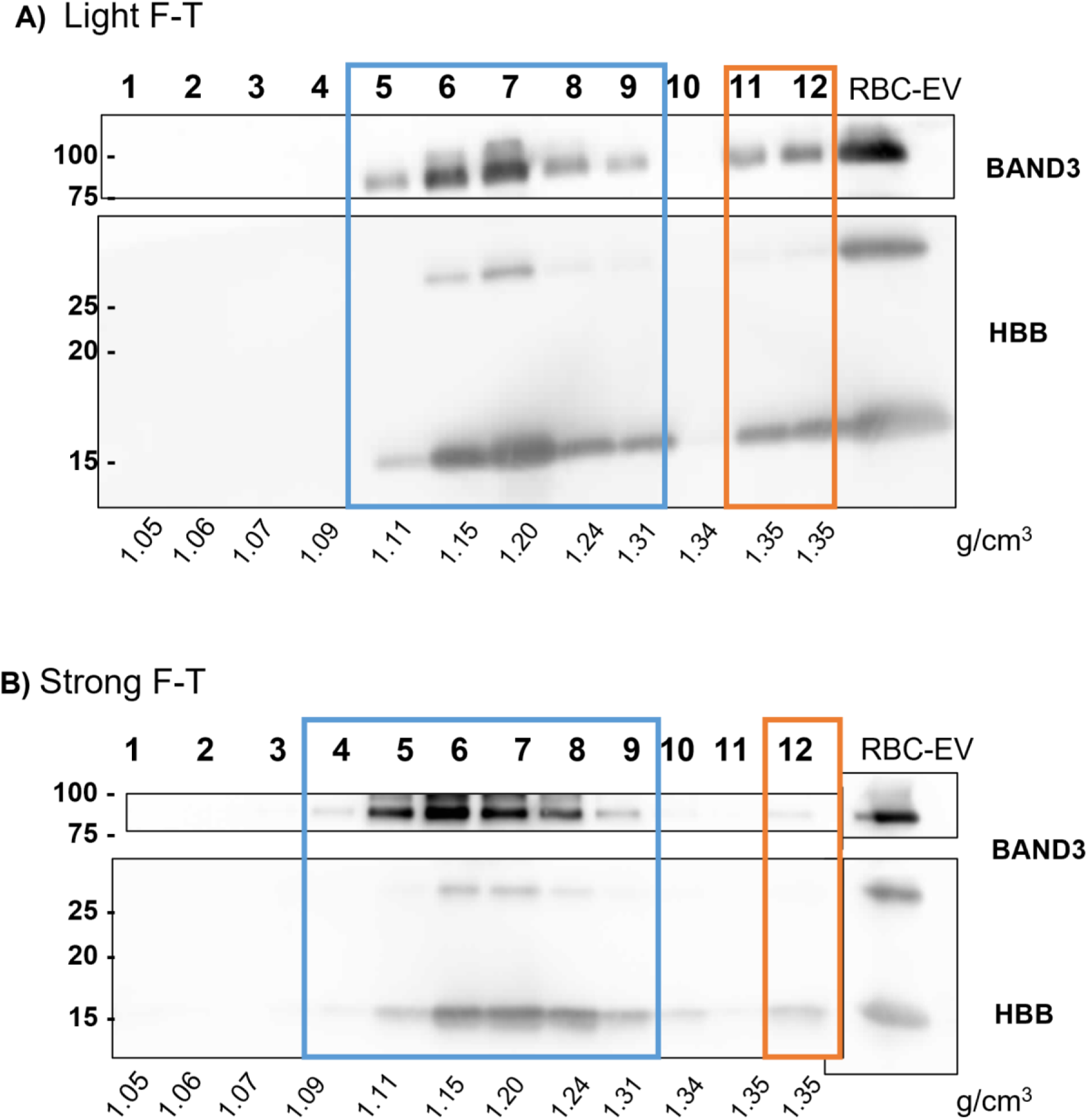
Gradient distribution of RBC-EVs after Freeze-Thaw cycles. Equal volume (15 μL) of each fraction both for Light Freeze-Thaw cycles (Light F-T, A), Strong Freeze-Thaw cycles (Strong F-T, B), 30 μg of RBC-EV homogenate (RBC-EV) samples were loaded. Samples were electrophoresed on SDS-PAGE gel (12% (Acrylamide/Bis-acrylamide) and analyzed for the antibodies described in the figures. Un-cropped WB are available in the Supplementary Information (Supplementary Fig.S6). Each fraction density is indicated in g/cm^3^. RBC-EV subpopulations with different gradient distributions are indicated in colored boxes: **A**) blue box fractions 5-9; orange fractions 11-12; **B**) blue box fractions 4-9; C) blue box 12.

These results suggest that both Light F-T and Strong F-T treatments reduce the heterogeneity of RBC-EV preparations in a “dose-dependent manner,” where the intensity of the treatment correlates with a greater degree of change. The observed trend indicates a shift in distribution that more closely resembles the pattern seen in the 6-month sample. This shift is also reflected in iNTA measurements. In the Light F-T samples, the relative concentration of EVs in the Upper subpopulation (contrast^1/3^ > 0.7) decreased from 70% to 65%, while the Lower subpopulation (contrast^1/3^ < 0.7) increased from 30% to 35%, though this difference was not statistically significant compared to Fresh samples (Fig. 5B, E, G). However, in the Strong F-T sample, the effect was more pronounced, with a statistically significant shift to 25% in the Upper subpopulation and 75% in the Lower subpopulation compared to Light F-T (Fig. 5C, F, G).

**Fig. 5.**
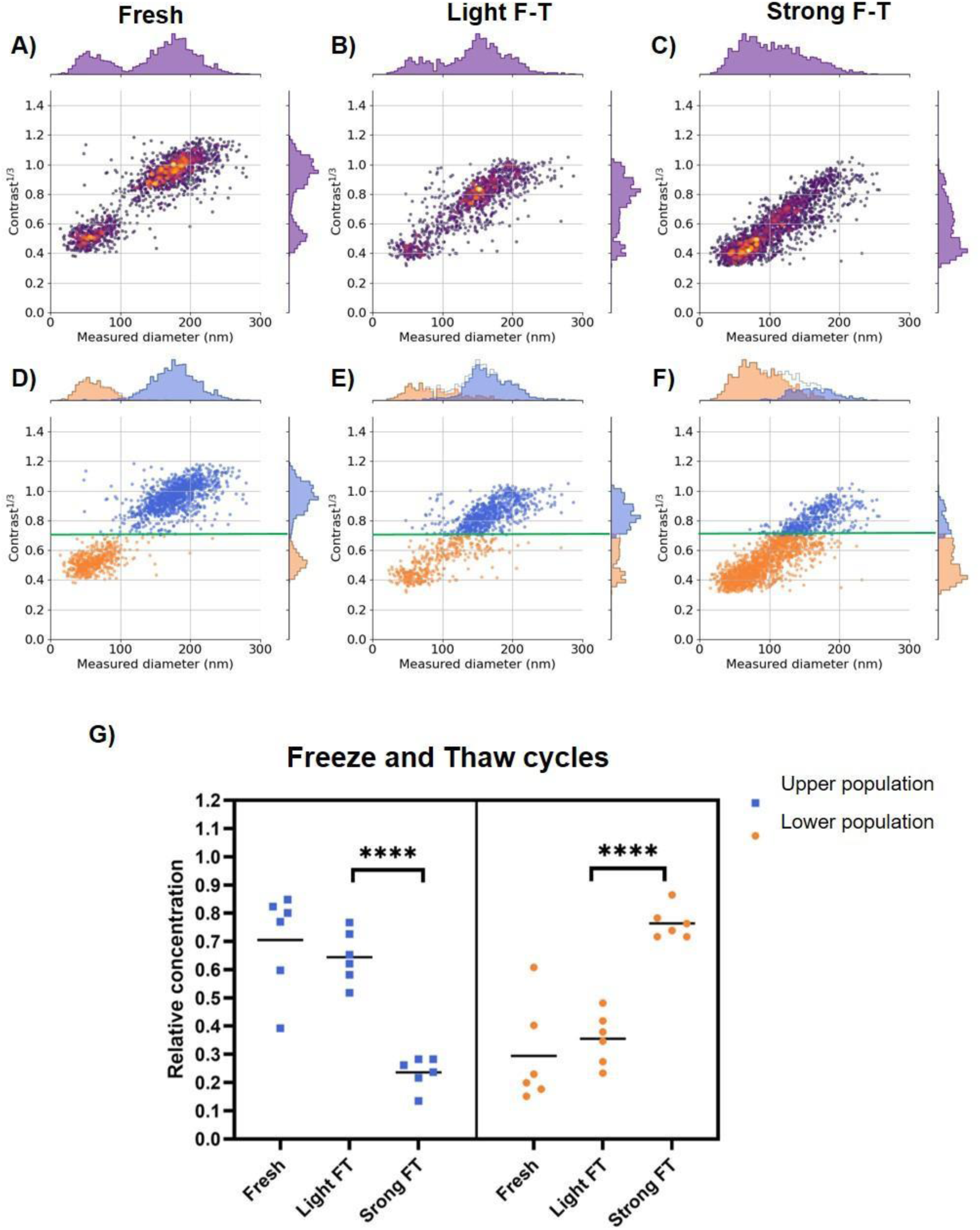
Size distribution and contrast evaluation of RBC-EVs after Freeze-Thaw cycles. Analysis by iNTA of Fresh (**A**), Light F-T (**B**), and Strong F-T (**C**) samples. Images represent size distribution (x-axis) and contrast (y-axis) of the nanoparticles. Nanoparticles subpopulations grouped for their contrast in Fresh (**D**), Light F-T (**E**), and Strong F-T (**F**) samples. The green line represents a contrast^1/3^ value of 0.7. Nanoparticle distribution determined for contrast^1/3^ ˃ 0.7 (blue dots, Upper population) or ˂ 0.7 (orange dots, Lower population). **G**) Relative Quantification of nanoparticles present in the Upper (contrast ^1/3^ ˃ 0.7) and Lower (contrast ^1/3^ ˂ 0.7) population for Fresh, 6-month, and 1-year samples. Statistical analysis using Student t-test: **** p value ˂ 0.0001.

Notably, iNTA data provided a quantitative confirmation of the changes observed in DSG distribution. The Strong F-T treatment altered the physico-chemical properties of the Fresh samples, making their density and contrast more similar to those of 6-month stored samples. The distribution between the Upper and Lower populations in the 6-month and the Strong F-T samples was not statistically different, with median relative concentrations of 20% and 25% in the upper subpopulation and 80% and 75% in the Lower subpopulation, respectively (Fig. 3G, Fig. 5G). We hypothesized that F-T cycles act as an “accelerated aging” method, reducing sample heterogeneity by mimicking the effects of long-term storage in a shorter time.

Accelerated aging is a strategy usually applied to a wide variety of samples: synthetic micro^61^ and nanoparticles^62^, organic^63^ and synthetic^64^ polymers^65^, plant seeds^66^, and pharmaceuticals^67^. Samples are subjected to different types of treatments^67^. The evaluation of the changes induced helps to determine the long-term effects of expected levels of stress within a shorter time^68^ and to select samples with determined characteristics and performances^66^.

Size distribution is a physicochemical characteristic that can be modified after different long-term storage conditions, both for EVs^69^ and synthetic nanoparticles^70^. In addition, accelerated aging can be applied to control this parameter in synthetic materials^62^.

Building on this, we used iNTA to investigate the differences in the size-distribution of nanoparticles in the five RBC-EVs samples: Fresh, 6-month, 1-year, Light F-T, and Strong F-T. In the Fresh sample (Fig.3A), two distinct subpopulations-below 100 nm (30-100 nm Fig 6A, B; purple histograms) and above 100 nm (Fig.6A, C; green histograms)-were easily distinguishable. Using these defined size ranges, we quantified the number of EVs in all samples.

**Fig. 6.**
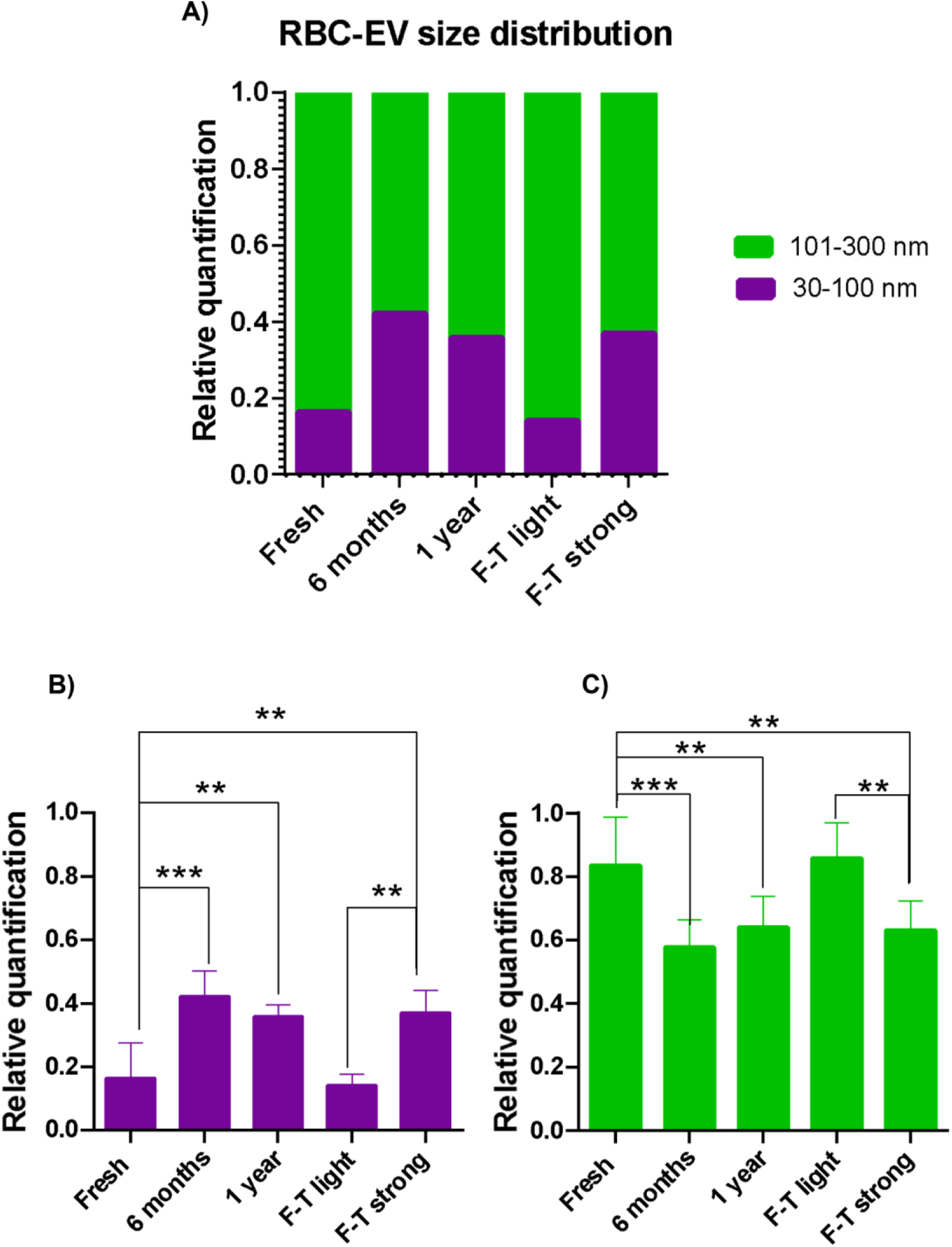
Size distribution evaluation of RBC-EVs after different storage time and F-T cycles. Characterization of RBC-EVs with iNTA. **A**) Relative quantification of nanoparticles with a diameter ranging from 30 to 100 nm (purple bar) and from 101 to 300 nm (green bar). **B**) Statistical analysis (ANOVA test) of changes in the size distribution for nanoparticles from 30 to 100 nm, **C**) Statistical analysis of nanoparticles from 101 to 300 nm. ** p value ≤ 0.01, *** p value ≤ 0.001

As shown in Fig. 6A, the Fresh sample has a relative concentration of 0.16 for particles below 100 nm (purple histogram) and 0.84 for particles above 100 nm (green histogram). After 6-month and 1-year of storage at -80 °C, the size distribution of RBC-EVs shifted significantly. Specifically, the relative concentration of the subpopulation below 100 nm increased to 0.42 in the 6-month sample and to 0.36 in the 1-year sample, both statistically significant compared to the Fresh sample (Fig. 6A,B). Consequently, the subpopulation above 100 nm decreased to 0.58 in the 6-month sample and 0.64 in the 1-year sample (Fig. 6C). These results indicate that long-term storage alters the relative size distribution of the RBC-EV nanoparticles, leading to an increased proportion of smaller particles.

This trend is further supported by the accelerated aging process, as the Strong F-T sample exhibited a statistically significant increase in the relative concentration of particles below 100 nm compared to the Fresh and Light F-T samples. Specifically, the proportion rose from 0.16 in the Fresh sample and 0.14 in the Light F-T sample to 0.37 in the Strong F-T sample (Fig. 6A, B). Taken together, these data demonstrate that after the Strong F-T treatment, RBC-EV physico-chemical characteristics resemble those of the sample stored 6 months at -80°C.

This effect has been described for liposome synthesis where extrusion, sonication, or ultrasound^71^ are used to break up multilamellar lipid vesicles and reduce liposome size^72^. As for density and contrast, we hypothesize that freezing processing could perturb RBC-EVs, and at room temperature, the vesicles tend to reform with a smaller size. Freezing typically leads to the formation of ice crystals, which can physically rupture cellular membranes or nanoparticles^73^, including vesicles like RBC-EVs. This mechanical stress could result in the breakdown of larger vesicles or a reduction in their size. Upon F-T processing, the two EV subpopulations might mix, reassembling in a more compact form, leading to a reduction in their size^74^. This process may be a stochastic one, as the phospholipid bilayer could reform spontaneously, potentially driven by amphiphilic nature and geometry, associate in an aqueous environment to form membranes^75^ rather than active molecular machinery (i.e. endosomal sorting complex required for transport (ESCRT) machinery for EV biogenesis)^76^ responsible for vesicle formation. Compared to their original state, this reorganization could result in smaller and less dense particles, originated by random encapsulation events^77^. Moreover, the results observed in this study are consistent with findings in other nanoparticle systems, including other types of extracellular vesicles (e.g. plasma EVs)^69,73^, where freezing and thawing can significantly alter size distribution and lead to a narrowing of the size range. The observed trend in RBC-EVs suggests that the size reduction and narrowing of size distribution over prolonged storage or freeze-thaw cycles might be a general phenomenon that could affect various types of EVs or lipid-based nanoparticles.

In conclusion, by measuring density, contrast^1/3^ and size distribution we showed that Strong F-T treatment produces RBC-EVs with properties resembling long-term stored samples. This approach may be useful for studying storage-related changes or for selecting samples that mimic long-term stored EVs.

### 3.4 Accelerated aging influences RBC-EV cellular uptake but not their surface protein activity

It has been demonstrated that biophysical and biochemical properties of biogenic^58,78^ and synthetic nanoparticles^79,80^ influence their biological activities. In order to analyze if the modifications induced in RBC-EV preparations after prolonged storage and accelerated aging could affect their interaction with cells and surface protein activity we analyzed their cellular uptake and enzymatic activity. To assess their interaction with cells we incubated equal particle numbers of fluorescently labelled RBC-EV from Fresh, 6-month, and Strong F-T on triple-negative breast cancer MDA-MA-231 cells for 30 min and 4h. Infact, we previously demonstrated a significant uptake of RBC-EV by this cell line^8^. Flow cytometry was performed to quantify the RBC-EV uptake. Analyses revealed that Fresh RBC-EVs were internalized by MDA-MB-231 cells to a higher extent than 6-month and Strong F-T RBC-EV both at shorter (30 min) and at extended (4h) time points. No significant differences between the uptake of EVs subjected to repeated 6-month and Strong F-T were highlighted, thus suggesting that the protocol including freeze-thaw cycles is effective in mimicking the prolonged storage at -80 °C (Fig 7A and 7B). Any unspecific fluorescent signal arising either from possibly increased cell autofluorescence due to EV internalization or to unbound MemGlow™ 488 fluorescent dye can be excluded: indeed, the signal from the aforementioned two experimental controls perfectly overlays the signal of the negative control, i.e., untreated cells (Supplementary Figure S7). It is worth noting that after 4 h incubation, the positivity of cells increases four times, regardless of the experimental condition (Fig. 7B). These results indicated that narrowing the heterogeneity of RBC-EVs can influence their interaction with target cells.

**Fig. 7.**
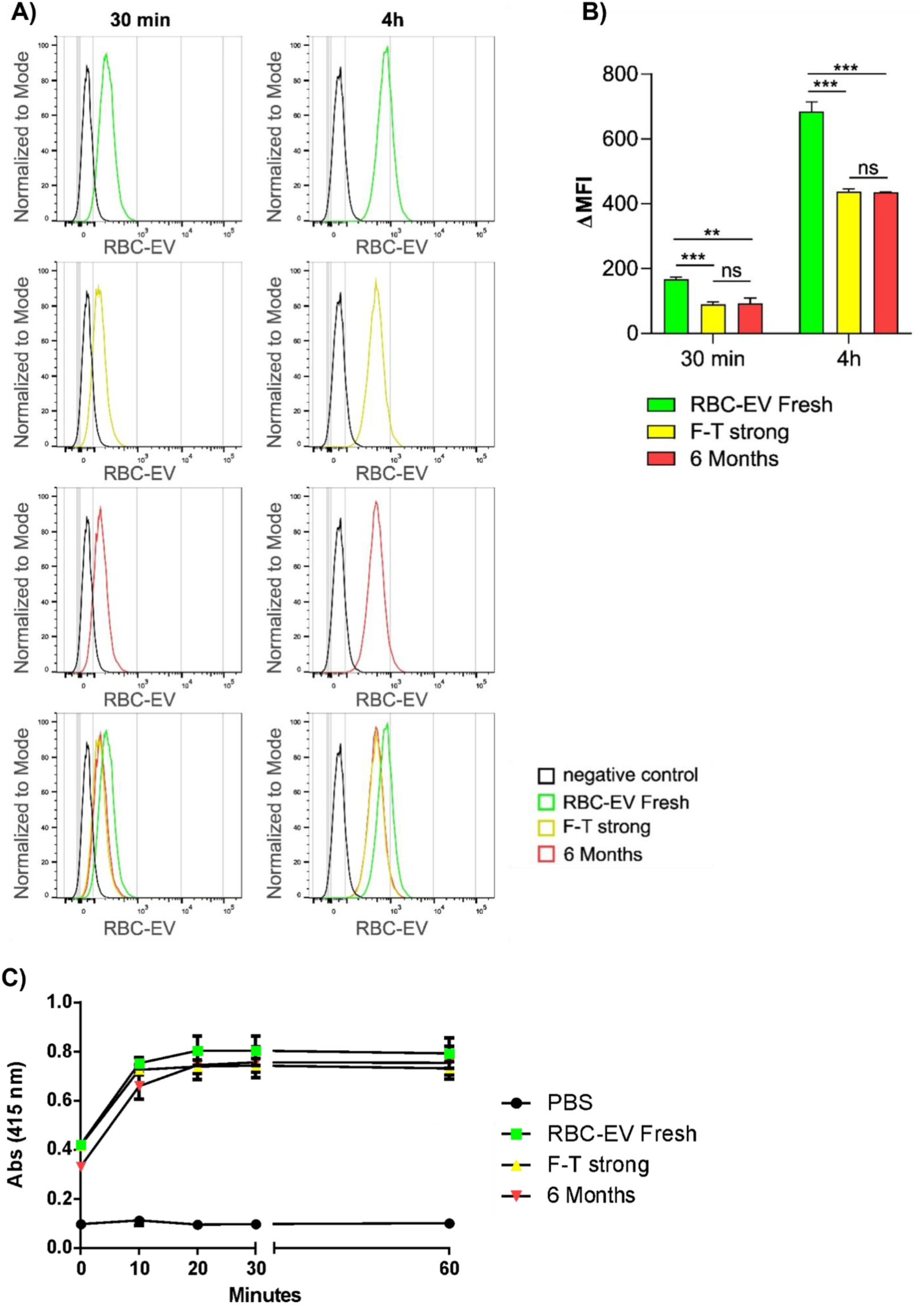
RBC-EV preparations biological activity. **A)** Flow cytometry analysis of RBC-EV uptake by MDA-MB-231 cells. Fresh (green), Strong F-T (yellow) and 6-month (red) samples have been stained with MemGlow™ 488 fluorescent dye and incubated with MDA-MB-231 cells for either 30 min or 4h. Cells have been detached with trypsin to obtain single-cell suspensions. Representative histograms are shown for both 30 min and 4h incubation. The fluorescence signal is compared to a negative control consisting of untreated cells. An overlay of all the experimental points is provided for both time points. **B**) Bars represent mean ± standard deviation of median fluorescence intensity (MFI) - MFI of negative control (ΔMFI), n=3, ** p<0.01, *** p<0.001, ns= not significant, Student’s t test. **C**) Acetylcholinesterase enzymatic activity measured after incubation of Fresh (green square), 6-month (red triangle down) and Strong F-T (yellow triangle up) samples with 1.25 mM acetylthiocholine and 0.1 mM 5,5’-dithio-bis(2-nitrobenzoic acid) in a final volume of 1 ml. Equal volume of PBS was used as negative control (black dots) The incubation was carried out at 37°C, and the change in absorbance at 415 nm was followed after 0, 10, 20, 30, and 60 min.

In addition, we wanted to verify if prolonged storage and F-T cycles affect the surface proteins present on the external membrane of RBC-EV. To test this, we analyzed the activity of acetylcholinesterase, a glycoprotein with enzymatic activity^81^ present on the surface of Red Blood cells and thus on RBC-EVs^82^, in Fresh, 6-months, and Strong F-T cycles samples. As shown in Fig.7C there were no significant differences in the enzymatic activity of the analyzed samples, indicating that the biological activity of enzymes present on the surface of the RBC-EVs are not altered neither by prolonged storage nor by Strong F-T cycles. This finding suggests two important points: 1) that reduced cellular uptake of stored EVs is likely due to factors other than surface enzyme degradation and 2) that the preservation of AChE activity highlights the robustness of the RBC-EV membrane’s functional integrity. However, while AChE activity was maintained, further studies should analyze other surface proteins for a comprehensive evaluation and comparison with other proteins that could be either transmembrane or part of the biomolecular corona^8,83,84^.

## 4. Conclusion

EVs are diverse nanoparticles with large heterogeneity in size and molecular composition. Although this heterogeneity provides high diagnostic value for liquid biopsy and confers many exploitable functions for therapeutic applications in cancer detection, wound healing and neurodegenerative and cardiovascular diseases, it has also impeded their clinical translation— hence heterogeneity acts as a double-edged sword^85^.

EV application as biomaterials for biological delivery systems may be affected by their complexity and this may pose a challenge for their manufacturing at large-scale^26^. Rationalizing this complexity in the context of the physico-chemical landscape will unlock possibilities in bioengineering^86^ and bio-nanoterapeutics towards personalized medicine.

In this study, we developed an “accelerated-aging” protocol based on freeze-thaw cycles that allows to obtain highly homogeneous RBC-EV preparations in terms of density, size distribution, and biological activity of RBC-EVs, able to mimic the biophysical characteristics of preparations stored for 6 months at -80°C. Our findings highlighted that long-term storage effectively reduced RBC-EV heterogeneity and led to a shift towards smaller and less dense particle subpopulation. Furthermore, our experiments showed that the reduced heterogeneity and smaller size of RBC-EVs, induced by either long-term storage or F-T cycles, impacted their cellular uptake in a similar way. This suggests that the subpopulation present in Fresh RBC-EVs that is able to give the highest uptake yield is the most unstable at the different storage conditions.

The described phenomena could be either due to extrinsic or intrinsic stochastic effects. Stochastic reactions can be simulated *in silico* by considering the number of molecules of a reacting system (e.g. the aqueous vesicle core), and calculating the probability that any possible reaction takes place accordingly. This opens a new perspective of using our model for “Synthetic Biology”^87,88^ experiments, representing concrete steps towards the identification of key active biomolecules/properties that determine EV mechanisms of action across different EV subtypes.

Overall, our results emphasize the importance of considering storage and processing conditions when preparing RBC-EVs for research or therapeutic applications, as these factors can significantly alter the properties and behavior of the vesicles. Additionally, the use of F-T cycles as an “accelerated aging” method offers a valuable tool for simulating long-term storage effects and studying the stability of RBC-EVs under various conditions.

This feature could be highly valuable in drug development, where standardizing RBC-EV preparations is essential to ensure reproducibility and therapeutic consistency. It also strongly supports the advancement of EVs as innovative biomaterials, both for cutting-edge research, such as surface protein engineering, and clinical applications, that demand stable and uniform preparations

## Author contributions

Conceptualization: LP, MR, AR; Data curation: LP, MR, VM, ST, SJ, AK, ELM, AM, AZ; Funding acquisition: PB, AR, LP; Investigation: LP, MR, ACB, SCG, PB, AR; Methodology: LP, MR, SJ, AK, VS Writing – original draft: LP writing review & editing: all authors.

## Acknowledgments

This work was supported by the Italian Ministry of Health (Ministero della Salute) through the Ricerca Finalizzata (RF) “Theory enhancing” grant: ID project RF-2021-12375279 and Project CN3 PNRR-Centro Nazionale sullo sviluppo di terapia genica e farmaci con tecnologia a RNA. The authors would like to thank the “Labion lab” of the IRCCS Fondazione Don Carlo Gnocchi ONLUS, Milan, Italy, for NTA measurements.

## Data availability

Data are available upon request.

## References

1. Théry C, Witwer KW, Aikawa E, Alcaraz MJ, Anderson JD, Andriantsitohaina R, et al. Minimal information for studies of extracellular vesicles 2018 (MISEV2018): a position statement of the International Society for Extracellular Vesicles and update of the MISEV2014 guidelines. J Extracell Vesicles. 2018 Dec 1;7(1):1535750.

2. Petraroia I, Ghidotti P, Bertolini G, Pontis F, Roz L, Balsamo M, et al. Extracellular vesicles from subjects with COPD modulate cancer initiating cells phenotype through HIF-1α shuttling. Cell Death Dis. 2023 Oct 14;14(10):681.

3. Raab-Traub N, Dittmer DP. Viral effects on the content and function of extracellular vesicles. Nat Rev Microbiol. 2017 Sept;15(9):559–72.

4. Kalluri R, LeBleu VS. The biology, function, and biomedical applications of exosomes. Science. 2020 Feb 7;367(6478):eaau6977.

5. Kumar SK, Sasidhar MV. Recent Trends in the Use of Small Extracellular Vesicles as Optimal Drug Delivery Vehicles in Oncology. Mol Pharm. 2023 Aug 7;20(8):3829–42.

6. Herrmann IK, Wood MJA, Fuhrmann G. Extracellular vesicles as a next-generation drug delivery platform. Nat Nanotechnol. 2021 July;16(7):748–59.

7. Pham CT, Zhang X, Lam A, Le MT. Red blood cell extracellular vesicles as robust carriers of RNA-based therapeutics. Cell Stress. 2018 Sept 10;2(9):239–41.

8. Musicò A, Zenatelli R, Romano M, Zendrini A, Alacqua S, Tassoni S, et al. Surface functionalization of extracellular vesicle nanoparticles with antibodies: a first study on the protein corona “variable.” Nanoscale Adv. 2023;5(18):4703–17.

9. Pham TC, Jayasinghe MK, Pham TT, Yang Y, Wei L, Usman WM, et al. Covalent conjugation of extracellular vesicles with peptides and nanobodies for targeted therapeutic delivery. J Extracell Vesicles. 2021 Feb;10(4):e12057.

10. Pham TT, Chen H, Nguyen PHD, Jayasinghe MK, Le AH, Le MT. Endosomal escape of nucleic acids from extracellular vesicles mediates functional therapeutic delivery. Pharmacol Res. 2023 Feb;188:106665.

11. Nguyen PHD, Jayasinghe MK, Le AH, Peng B, Le MTN. Advances in Drug Delivery Systems Based on Red Blood Cells and Their Membrane-Derived Nanoparticles. ACS Nano. 2023 Mar 28;17(6):5187–210.

12. Repsold L, Joubert AM. Eryptosis: An Erythrocyte’s Suicidal Type of Cell Death. BioMed Res Int. 2018;2018:9405617.

13. Dodson, R.A., Hinds, T.R. and Vincenzi, F.F. (1987) Effects of Calcium and A23187 on Deformability and Volume of Human Red Blood Cells. Blood Cells, 12, 555–564. - References - Scientific Research Publishing [Internet]. [cited 2025 Aug 6]. Available from: https://www.scirp.org/reference/referencespapers?referenceid=2565112

14. Hasse S, Duchez AC, Fortin P, Boilard E, Bourgoin SG. Interplay between LPA2 and LPA3 in LPA-mediated phosphatidylserine cell surface exposure and extracellular vesicles release by erythrocytes. Biochem Pharmacol. 2021 Oct 1;192:114667.

15. Nguyen DB, Thuy Ly TB, Wesseling MC, Hittinger M, Torge A, Devitt A, et al. Characterization of microvesicles released from human red blood cells. Cell Physiol Biochem. 2016 Apr 3;38(3):1085–99.

16. Ebeyer-Masotta M, Eichhorn T, Fischer MB, Weber V. Impact of production methods and storage conditions on extracellular vesicles in packed red blood cells and platelet concentrates. Transfus Apher Sci. 2024 Apr 1;63(2):103891.

17. Gamonet C, Desmarets M, Mourey G, Biichle S, Aupet S, Laheurte C, et al. Processing methods and storage duration impact extracellular vesicle counts in red blood cell units. Blood Adv. 2020 Nov 10;4(21):5527–39.

18. Usman WM, Pham TC, Kwok YY, Vu LT, Ma V, Peng B, et al. Efficient RNA drug delivery using red blood cell extracellular vesicles. Nat Commun. 2018 Dec;9(1):2359.

19. Prudent M, Crettaz D, Delobel J, Seghatchian J, Tissot JD, Lion N. Differences between calcium-stimulated and storage-induced erythrocyte-derived microvesicles. Transfus Apher Sci. 2015 Oct 1;53(2):153–8.

20. Salzer U, Zhu R, Luten M, Isobe H, Pastushenko V, Perkmann T, et al. Vesicles generated during storage of red cells are rich in the lipid raft marker stomatin. Transfusion (Paris). 2008;48(3):451–62.

21. de Oliveira GP, Welsh JA, Pinckney B, Palu CC, Lu S, Zimmerman A, et al. Human red blood cells release microvesicles with distinct sizes and protein composition that alter neutrophil phagocytosis. J Extracell Biol. 2023 Oct 25;2(11):e107.

22. Kusuma GD, Barabadi M, Tan JL, Morton DAV, Frith JE, Lim R. To Protect and to Preserve: Novel Preservation Strategies for Extracellular Vesicles. Front Pharmacol [Internet]. 2018 Oct 29 [cited 2025 Aug 6];9. Available from: https://www.frontiersin.org/journals/pharmacology/articles/10.3389/fphar.2018.01199/full

23. Almizraq RJ, Seghatchian J, Holovati JL, Acker JP. Extracellular vesicle characteristics in stored red blood cell concentrates are influenced by the method of detection. Transfus Apher Sci. 2017 Apr 1;56(2):254– 60.

24. Bebesi T, Kitka D, Gaál A, Szigyártó IC, Deák R, Beke-Somfai T, et al. Storage conditions determine the characteristics of red blood cell derived extracellular vesicles. Sci Rep. 2022 Jan 19;12(1):977.

25. Straat M, Böing AN, Tuip-De Boer A, Nieuwland R, Juffermans NP. Extracellular Vesicles from Red Blood Cell Products Induce a Strong Pro-Inflammatory Host Response, Dependent on Both Numbers and Storage Duration. Transfus Med Hemotherapy Off Organ Dtsch Ges Transfusionsmedizin Immunhamatologie. 2016 July;43(4):302–5.

26. Paolini L, Monguió-Tortajada M, Costa M, Antenucci F, Barilani M, Clos-Sansalvador M, et al. Large-scale production of extracellular vesicles: Report on the “massivEVs” ISEV workshop. J Extracell Biol. 2022 Oct 25;1(10):e63.

27. Nelson BC, Maragh S, Ghiran IC, Jones JC, DeRose PC, Elsheikh E, et al. Measurement and standardization challenges for extracellular vesicle therapeutic delivery vectors. Nanomed. 2020 Sept;15(22):2149–70.

28. ICH Q1A(R2) Stability Testing of New Drug Substances and Products [Internet]. Food and Drug Administration; 2003 Nov [cited 2024 Oct 31]. Available from: https://www.fda.gov/regulatory-information/search-fda-guidance-documents/q1ar2-stability-testing-new-drug-substances-and-products

29. ICH Q1A (R2) Stability testing of new drug substances and drug products - Scientific guideline [Internet]. European Medicines Agency (EMA); 2003 Feb [cited 2024 Oct 31]. Available from: https://www.ema.europa.eu/en/ich-q1a-r2-stability-testing-new-drug-substances-drug-products-scientific-guideline#current-effective-version-8899

30. Saugstad JA, Lusardi TA, Van Keuren-Jensen KR, Phillips JI, Lind B, Harrington CA, et al. Analysis of extracellular RNA in cerebrospinal fluid. J Extracell Vesicles. 2017;6(1):1317577.

31. Tian Y, Gong M, Hu Y, Liu H, Zhang W, Zhang M, et al. Quality and efficiency assessment of six extracellular vesicle isolation methods by nano-flow cytometry. J Extracell Vesicles. 2020 Jan 1;9(1):1697028.

32. Gori A, Romanato A, Bergamaschi G, Strada A, Gagni P, Frigerio R, et al. Membrane-binding peptides for extracellular vesicles on-chip analysis. J Extracell Vesicles. 2020;9(1):1751428.

33. Ridolfi A, Brucale M, Montis C, Caselli L, Paolini L, Borup A, et al. AFM-Based High-Throughput Nanomechanical Screening of Single Extracellular Vesicles. Anal Chem. 2020 Aug 4;92(15):10274–82.

34. Raposo G, Stoorvogel W. Extracellular vesicles: exosomes, microvesicles, and friends. J Cell Biol. 2013 Feb 18;200(4):373–83.

35. Carlomagno C, Giannasi C, Niada S, Bedoni M, Gualerzi A, Brini AT. Raman Fingerprint of Extracellular Vesicles and Conditioned Media for the Reproducibility Assessment of Cell-Free Therapeutics. Front Bioeng Biotechnol [Internet]. 2021 Apr 13 [cited 2025 Aug 6];9. Available from: https://www.frontiersin.org/journals/bioengineering-and-biotechnology/articles/10.3389/fbioe.2021.640617/full

36. Paolini L, Federici S, Consoli G, Arceri D, Radeghieri A, Alessandri I, et al. Fourier-transform Infrared (FT-IR) spectroscopy fingerprints subpopulations of extracellular vesicles of different sizes and cellular origin. J Extracell Vesicles. 2020 Sept;9(1):1741174.

37. Koponen A, Kerkelä E, Rojalin T, Lázaro-Ibáñez E, Suutari T, Saari HO, et al. Label-free characterization and real-time monitoring of cell uptake of extracellular vesicles. Biosens Bioelectron. 2020 Nov 15;168:112510.

38. Kashkanova AD, Blessing M, Gemeinhardt A, Soulat D, Sandoghdar V. Precision size and refractive index analysis of weakly scattering nanoparticles in polydispersions. Nat Methods. 2022 May;19(5):586–93.

39. Kashkanova AD, Blessing M, Reischke M, Baur JO, Baur AS, Sandoghdar V, et al. Label-free discrimination of extracellular vesicles from large lipoproteins. J Extracell Vesicles. 2023 Aug;12(8):e12348.

40. Van Deun J, Mestdagh P, Agostinis P, Akay Ö, Anand S, Anckaert J, et al. EV-TRACK: transparent reporting and centralizing knowledge in extracellular vesicle research. Nat Methods. 2017 Mar;14(3):228–32.

41. Grossi I, Radeghieri A, Paolini L, Porrini V, Pilotto A, Padovani A, et al. MicroRNA–34a–5p expression in the plasma and in its extracellular vesicle fractions in subjects with Parkinson’s disease: An exploratory study. Int J Mol Med. 2020 Dec 2;47(2):533–46.

42. Radeghieri A, Alacqua S, Zendrini A, Previcini V, Todaro F, Martini G, et al. Active antithrombin glycoforms are selectively physiosorbed on plasma extracellular vesicles. J Extracell Biol. 2022 Sept;1(9):e57.

43. Paolini L, Radeghieri A, Civini S, Caimi L, Ricotta D. The Epsilon Hinge-Ear Region Regulates Membrane Localization of the AP-4 Complex. Traffic. 2011 Nov;12(11):1604–19.

44. Di Noto G, Cimpoies E, Dossi A, Paolini L, Radeghieri A, Caimi L, et al. Polyclonal versus monoclonal immunoglobulin-free light chains quantification. Ann Clin Biochem Int J Lab Med. 2015 May;52(3):327– 36.

45. Alvisi G, Paolini L, Contarini A, Zambarda C, Di Antonio V, Colosini A, et al. Intersectin goes nuclear: secret life of an endocytic protein. Biochem J. 2018 Apr 30;475(8):1455–72.

46. Salvi A, Vezzoli M, Busatto S, Paolini L, Faranda T, Abeni E, et al. Analysis of a nanoparticle–enriched fraction of plasma reveals miRNA candidates for Down syndrome pathogenesis. Int J Mol Med [Internet]. 2019 Apr 9 [cited 2025 Aug 6]; Available from: http://www.spandidos-publications.com/10.3892/ijmm.2019.4158

47. Shah JS, Soon PS, Marsh DJ. Comparison of Methodologies to Detect Low Levels of Hemolysis in Serum for Accurate Assessment of Serum microRNAs. Janigro D, editor. PLOS ONE. 2016 Apr 7;11(4):e0153200.

48. Maiolo D, Paolini L, Di Noto G, Zendrini A, Berti D, Bergese P, et al. Colorimetric Nanoplasmonic Assay To Determine Purity and Titrate Extracellular Vesicles. Anal Chem. 2015 Apr 21;87(8):4168–76.

49. Zendrini A, Paolini L, Busatto S, Radeghieri A, Romano M, Wauben MHM, et al. Augmented COlorimetric NANoplasmonic (CONAN) Method for Grading Purity and Determine Concentration of EV Microliter Volume Solutions. Front Bioeng Biotechnol. 2020 Feb 12;7:452.

50. Borup A, Boysen AT, Ridolfi A, Brucale M, Valle F, Paolini L, et al. Comparison of separation methods for immunomodulatory extracellular vesicles from helminths. J Extracell Biol. 2022 May;1(5):e41.

51. Kashkanova AD, Albrecht D, Küppers M, Blessing M, Sandoghdar V. Measuring Concentration of Nanoparticles in Polydisperse Mixtures Using Interferometric Nanoparticle Tracking Analysis. ACS Nano. 2024 July 23;18(29):19161–8.

52. Ho IK, Ellman GL. TRITON SOLUBILIZED ACETYLCHOLINESTERASE OF BRAIN. J Neurochem. 1969 Nov;16(11):1505–13.

53. Savina A, Vidal M, Colombo MI. The exosome pathway in K562 cells is regulated by Rab11. J Cell Sci. 2002 June 15;115(12):2505–15.

54. Welsh JA, Goberdhan DCI, O’Driscoll L, Buzas EI, Blenkiron C, Bussolati B, et al. Minimal information for studies of extracellular vesicles (MISEV2023): From basic to advanced approaches. J Extracell Vesicles. 2024 Feb;13(2):e12404.

55. Lucien F, Gustafson D, Lenassi M, Li B, Teske JJ, Boilard E, et al. MIBlood-EV: Minimal information to enhance the quality and reproducibility of blood extracellular vesicle research. J Extracell Vesicles. 2023 Dec;12(12):e12385.

56. Gori A, Frigerio R, Gagni P, Burrello J, Panella S, Raimondi A, et al. Addressing Heterogeneity in Direct Analysis of Extracellular Vesicles and Their Analogs by Membrane Sensing Peptides as Pan-Vesicular Affinity Probes. Adv Sci. 2024;11(29):2400533.

57. Musicò A, Zendrini A, Gimenez Reyes S, Mangolini V, Paolini L, Romano M, et al. Extracellular vesicles of different cellular origin feature distinct biomolecular corona dynamics. Nanoscale Horiz. 2025;10(1):104–12.

58. Caponnetto F, Manini I, Skrap M, Palmai-Pallag T, Di Loreto C, Beltrami AP, et al. Size-dependent cellular uptake of exosomes. Nanomedicine Nanotechnol Biol Med. 2017 Apr 1;13(3):1011–20.

59. Mangolini V, Radeghieri A, Piva S, Cattaneo S, Brucale M, Valle F, et al. Universal protocol to separate and compare Extracellular Vesicles from human plasma and skeletal muscle biopsy [Internet]. bioRxiv; 2025 [cited 2025 Aug 7]. p. 2024.02.19.580950. Available from: https://www.biorxiv.org/content/10.1101/2024.02.19.580950v2

60. Huang YX, Wu ZJ, Huang BT, Luo M. Pathway and Mechanism of pH Dependent Human Hemoglobin Tetramer-Dimer-Monomer Dissociations. PLOS ONE. 2013 Nov 28;8(11):e81708.

61. Cheng X, Wang S, Zhang X, Iqbal MS, Yang Z, Xi Y, et al. Accelerated aging behavior of degradable and non-degradable microplastics via advanced oxidation and their adsorption characteristics towards tetracycline. Ecotoxicol Environ Saf. 2024 Oct 1;284:116864.

62. Shuklov IA, Toknova VF, Lizunova AA, Razumov VF. Controlled aging of PbS colloidal quantum dots under mild conditions. Mater Today Chem. 2020 Dec 1;18:100357.

63. Calvini P, Gorassini A. On the Rate of Paper Degradation: Lessons From the Past. Restaurator [Internet]. 2006 Jan 19 [cited 2025 Aug 7];27(4). Available from: https://www.degruyter.com/document/doi/10.1515/REST.2006.275/html

64. Fambri L, Caria R, Atzori F, Ceccato R, Lorenzi D. Controlled Aging and Degradation of Selected Plastics in Marine Environment: 12 Months of Follow-up. In: Cocca M, Di Pace E, Errico ME, Gentile G, Montarsolo A, Mossotti R, et al., editors. Proceedings of the 2nd International Conference on Microplastic Pollution in the Mediterranean Sea. Cham: Springer International Publishing; 2020. p. 89– 100.

65. Kockott D. New method for accelerated testing of the aging behavior of polymeric materials as a function of radiation and temperature. Polym Test. 2022 June 1;110:107550.

66. Fenollosa E, Jené L, Munné-Bosch S. A rapid and sensitive method to assess seed longevity through accelerated aging in an invasive plant species. Plant Methods. 2020 May 8;16(1):64.

67. Waterman KC, Adami RC. Accelerated aging: prediction of chemical stability of pharmaceuticals. Int J Pharm. 2005 Apr 11;293(1–2):101–25.

68. Zou X, Uesaka T, Gurnagul N. Prediction of paper permanence by accelerated aging I. Kinetic analysis of the aging process. Cellulose. 1996 Dec;3(1):243–67.

69. Gelibter S, Marostica G, Mandelli A, Siciliani S, Podini P, Finardi A, et al. The impact of storage on extracellular vesicles: A systematic study. J Extracell Vesicles. 2022;11(2):e12162.

70. Gubicza J, Lábár JL, Quynh LM, Nam NH, Luong NH. Evolution of size and shape of gold nanoparticles during long-time aging. Mater Chem Phys. 2013 Mar 15;138(2):449–53.

71. Huang X, Caddell R, Yu B, Xu S, Theobald B, Lee LJ, et al. Ultrasound-enhanced microfluidic synthesis of liposomes. Anticancer Res. 2010 Feb;30(2):463–6.

72. Yun JS, Hwangbo SA, Jeong YG. Preparation of Uniform Nano Liposomes Using Focused Ultrasonic Technology. Nanomaterials. 2023 Jan;13(19):2618.

73. Sivanantham A, Jin Y. Impact of Storage Conditions on EV Integrity/Surface Markers and Cargos. Life. 2022 May 7;12(5):697.

74. Grosfils P, Losada-Pérez P. Kinetic control of liposome size by direct lipid transfer. J Colloid Interface Sci. 2023 Dec 15;652:1381–93.

75. Svetina S, Zeks B. Shape behavior of lipid vesicles as the basis of some cellular processes. Anat Rec. 2002 Nov 1;268(3):215–25.

76. Maas SLN, Breakefield XO, Weaver AM. Extracellular Vesicles: Unique Intercellular Delivery Vehicles. Trends Cell Biol. 2017 Mar;27(3):172–88.

77. Altamura E, Carrara P, D’Angelo F, Mavelli F, Stano P. Extrinsic stochastic factors (solute partition) in gene expression inside lipid vesicles and lipid-stabilized water-in-oil droplets: a review. Synth Biol. 2018 Jan 1;3(1):ysy011.

78. Wang X, Zhang Z, Qi Y, Zhang Z, Zhang Y, Meng K, et al. Study of the uptake mechanism of two small extracellular vesicle subtypes by granulosa cells. Anim Reprod Sci. 2024 Nov 1;270:107576.

79. Andar AU, Hood RR, Vreeland WN, Devoe DL, Swaan PW. Microfluidic preparation of liposomes to determine particle size influence on cellular uptake mechanisms. Pharm Res. 2014 Feb;31(2):401–13.

80. Jurney P, Agarwal R, Singh V, Choi D, Roy K, Sreenivasan SV, et al. Unique size and shape-dependent uptake behaviors of non-spherical nanoparticles by endothelial cells due to a shearing flow. J Control Release Off J Control Release Soc. 2017 Jan 10;245:170–6.

81. Daniels G. Functions of red cell surface proteins. Vox Sang. 2007 Nov;93(4):331–40.

82. Liao Z, Jaular LM, Soueidi E, Jouve M, Muth DC, Schøyen TH, et al. Acetylcholinesterase is not a generic marker of extracellular vesicles. J Extracell Vesicles. 2019 Dec;8(1):1628592.

83. Tóth EÁ, Turiák L, Visnovitz T, Cserép C, Mázló A, Sódar BW, et al. Formation of a protein corona on the surface of extracellular vesicles in blood plasma. J Extracell Vesicles [Internet]. 2021 Sept [cited 2022 July 25];10(11). Available from: https://onlinelibrary.wiley.com/doi/10.1002/jev2.12140

84. Tassoni S, Bergese P, Radeghieri A. The extracellular vesicle biomolecular corona: current insights and diagnostic potential. Nanomed. 2025 Aug;20(16):2013–21.

85. Carney RP, Mizenko RR, Bozkurt BT, Lowe N, Henson T, Arizzi A, et al. Harnessing extracellular vesicle heterogeneity for diagnostic and therapeutic applications. Nat Nanotechnol. 2025 Jan;20(1):14–25.

86. Manno M, Bongiovanni A, Margolis L, Bergese P, Arosio P. The physico-chemical landscape of extracellular vesicles. Nat Rev Bioeng. 2024 Nov 12;3(1):68–82.

87. Voigt CA. Synthetic biology 2020–2030: six commercially-available products that are changing our world. Nat Commun. 2020 Dec 11;11(1):6379.

88. Garner KL. Principles of synthetic biology. Essays Biochem. 2021 Nov 2;65(5):791–811.

