## Supplementary Info for "Red blood Cell-derived Extracellular Vesicles as biomaterials: the opportunity of freezing-induced accelerated aging"

d Dept. Molecular and Translational Medicine (DMMT), University of Brescia, Brescia, Italy

e IRCCS Fondazione Don Carlo Gnocchi ONLUS, Milano, Italy.

f Max-Planck-Institut für die Physik des Lichts, 91058 Erlangen, Germany

g Max-Planck-Zentrum für Physik und Medizin, 91058 Erlangen, Germany

h "Angelo Nocivelli" Institute for Molecular Medicine, ASST Spedali Civili, 25123, Brescia, Italy

i Department Physik, Friedrich-Alexander Universität Erlangen, Nürnberg, 91058 Erlangen, Germany

l Laboratory of Stem Cells, U.O.C. of Immunohaematology and Transfusion Medicine, Santo Spirito Hospital, Pescara, Italy

m Section of Medical Genetics and Cytogenetics, ASST Spedali Civili of Brescia, 25123 Brescia, Italy

n National Center for Gene Therapy and Drugs based on RNA Technology – CN3, Brescia, Italy

* These authors equally contributed to the work.

§ Corresponding author

### Supplementary Info

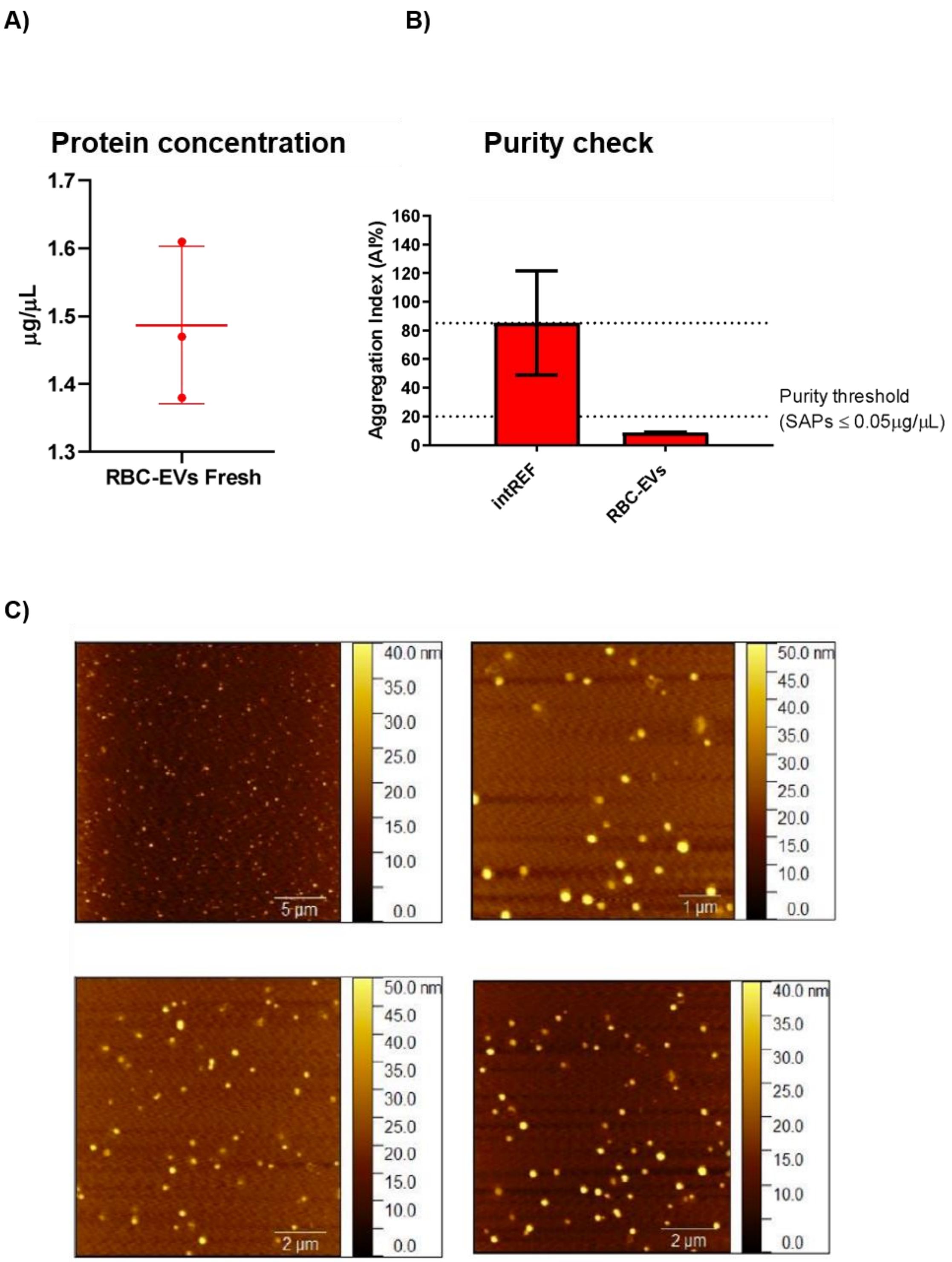


**Figure S1.** RBC-EV Fresh characterization. A) Protein concentration of Fresh RBC-EV calculated by Bradford assay. B) Purity check tested with the CONAN assay. IntREF: Internal reference control, aggregation index (AI) of AuNPs without Evs; RBC-EVs: AI of AuNPs after interaction with Fresh RBC-

EV preparation. Purity determined with AI under 20% indicating a contamination of Single and Aggregated Proteina (SAPs) below 0.05 ug/ul. (C) Representative images of Fresh RBC-EV sample by Atomic Force Microscopy (AFM) at different magnifications.

A)

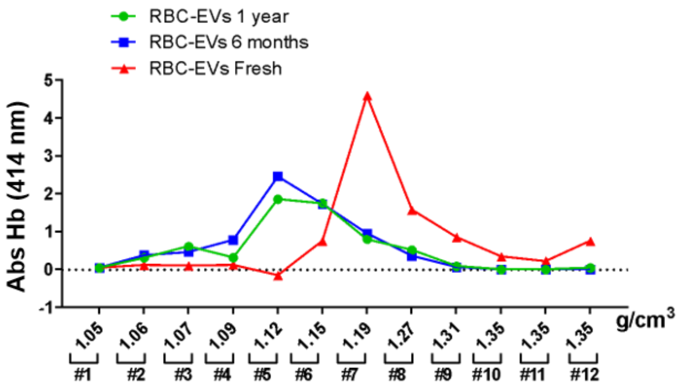


B)

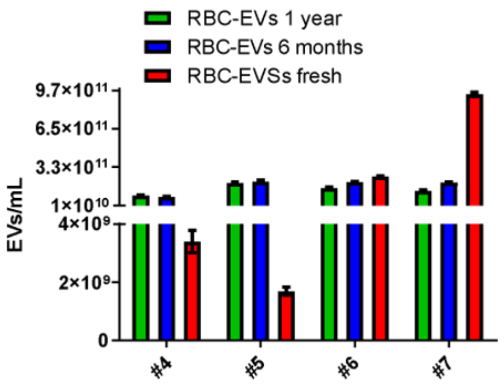


C)

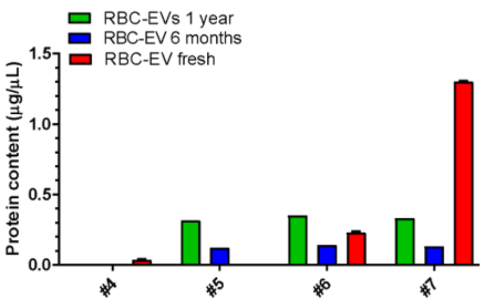


**Figure S2. Analysis of the RBC-EV Fresh, 6-month, 1-year samples after DSG processing. A) Hemoglobin distribution.** Hemoglobin distribution along DSG 12 fractions was checked determining hemoglobin absorbance at 414 nm for Fresh (red), 6-month (blue), 1-year (green) samples. C) Nanoparticle concentration from fraction 4 to fraction 7. RBC-EV concentration determined by Nanoparticle Tracking Analysis (NTA) for Fresh (red), 6 months (blue), 1-year (green) samples. D) Protein concentration from fraction 4 to fraction 7. RBC-EV protein concentration determined by Bicinchoninic acid (BCA) assay for Fresh (red), 6 month (blue), 1-year (green) samples.

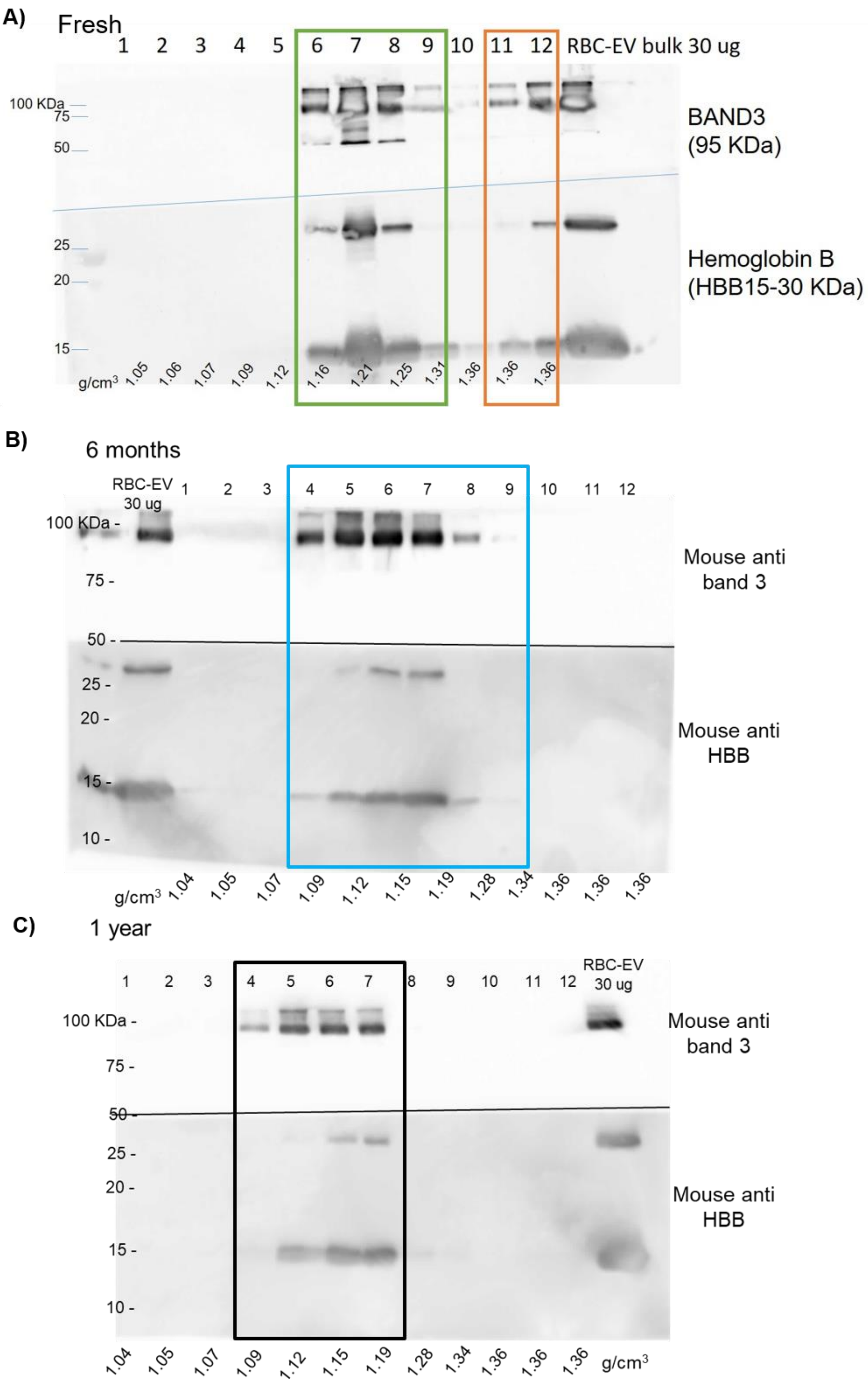


**Figure S3**. **Uncropped Western Blot images of gradient distribution of RBC-EVs after different storage conditions. A)** Fresh RBC-EVs**; B)** RBC-EVs stored for 6 months at -80; **C)** RBC-Evs stored for 1-year at -80°C**.**

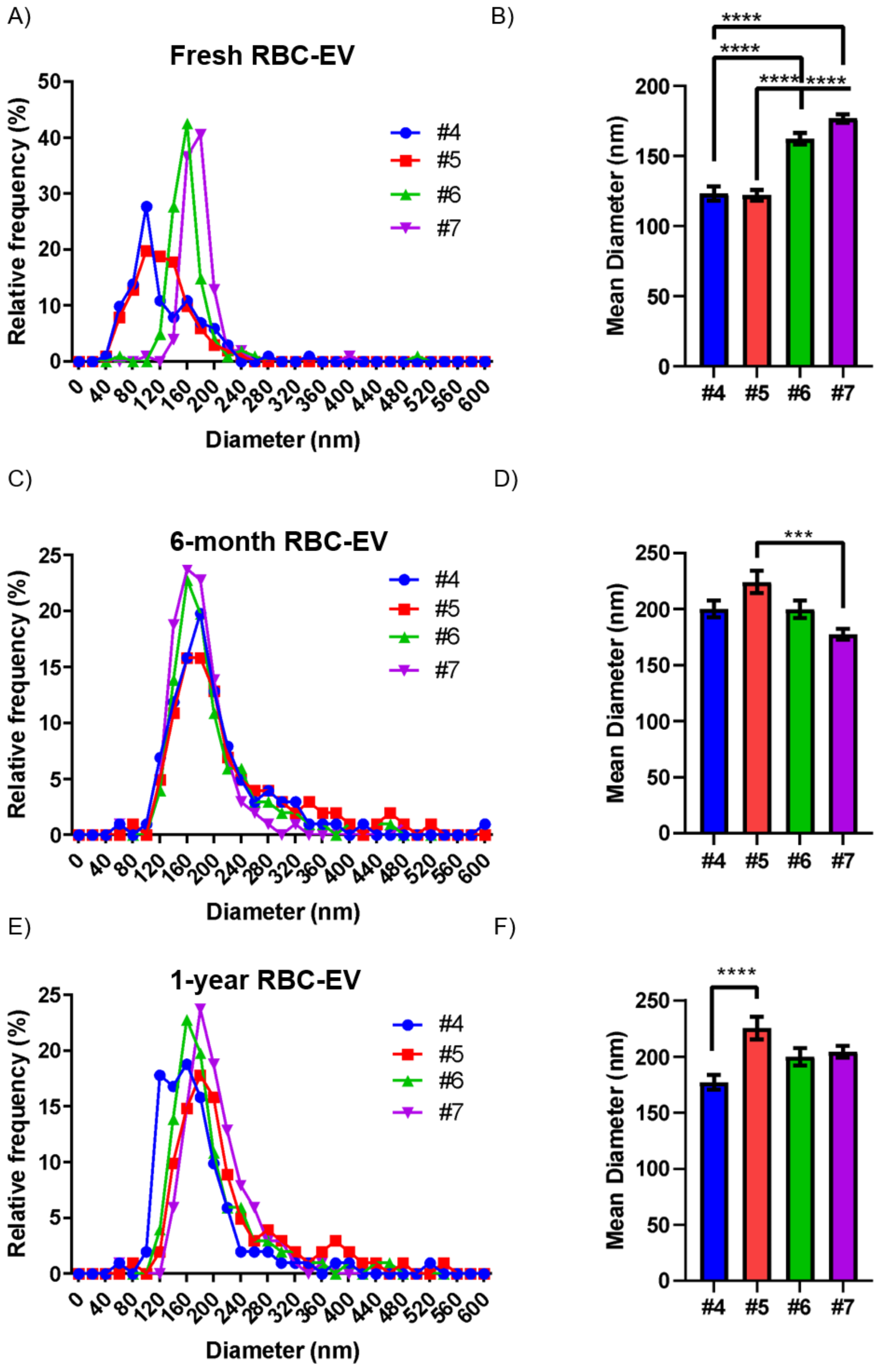


**Figure S4. Size distribution of RBC-EVs storaged in different conditions determined by NTA. A)** Size distribution nanoparticles in DSG fractions from 4 to 7 for Fresh RBC-Evs, stored at +4°C **B)** Size distribution of DSG fractions from 4 to 7 of 6-months RBC-Evs, stored at -80°C for 6 months **C)** Size distribution of DSG fractions from 4 to 7 of 1-year RBC-Evs, stored at -80°C for 1 year.

**A)**

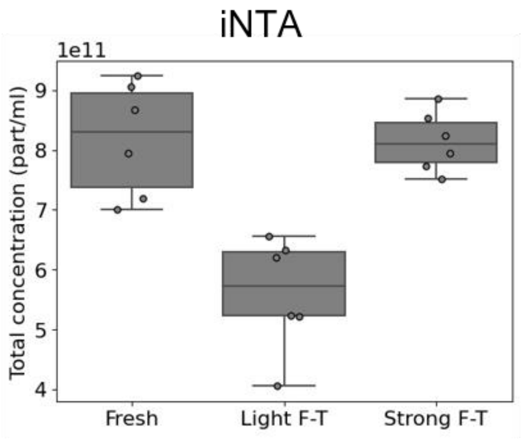


**B)**

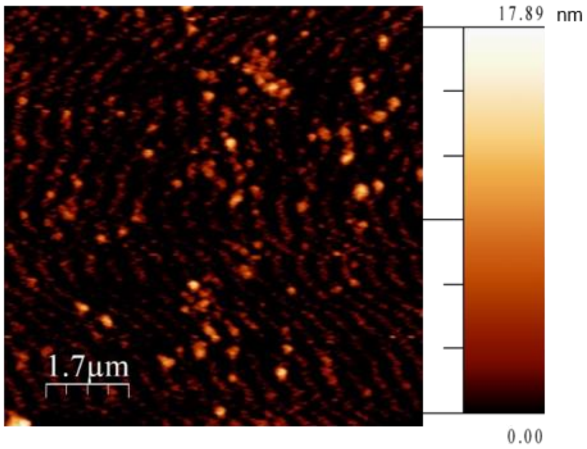


DSG fraction 4 of RBC-EV Strong F-T

**C)**

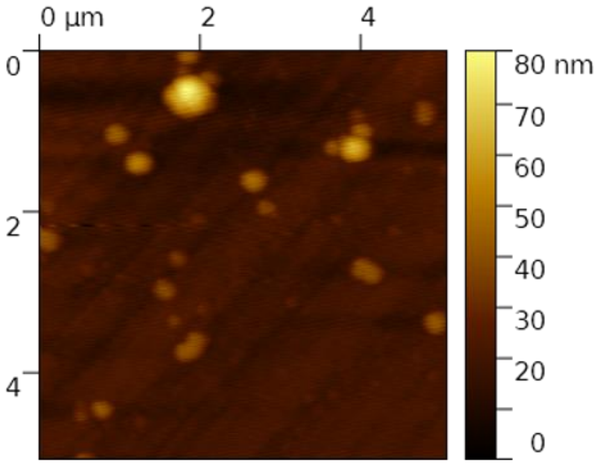


DSG fraction 7 of RBC-EV Strong F-T

**Figure S5: A)** Particle concentration mesured by iNTA in Fresh sample and after Light and Strong Freeze and thaw processing. B) AFM imaging of fraction 4 of DSG of Strong F-T sample. C) AFM imaging of fraction 7 of DSG of Strong F-T sample

A)

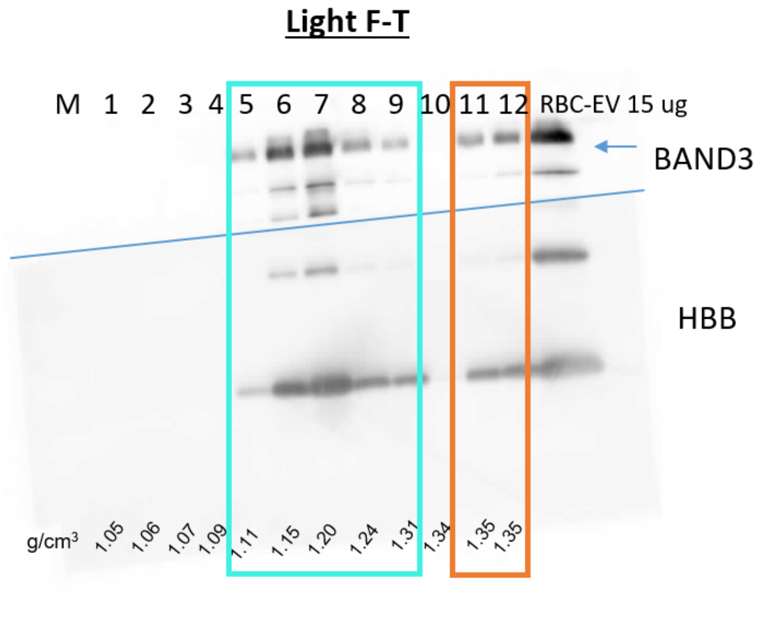


B)

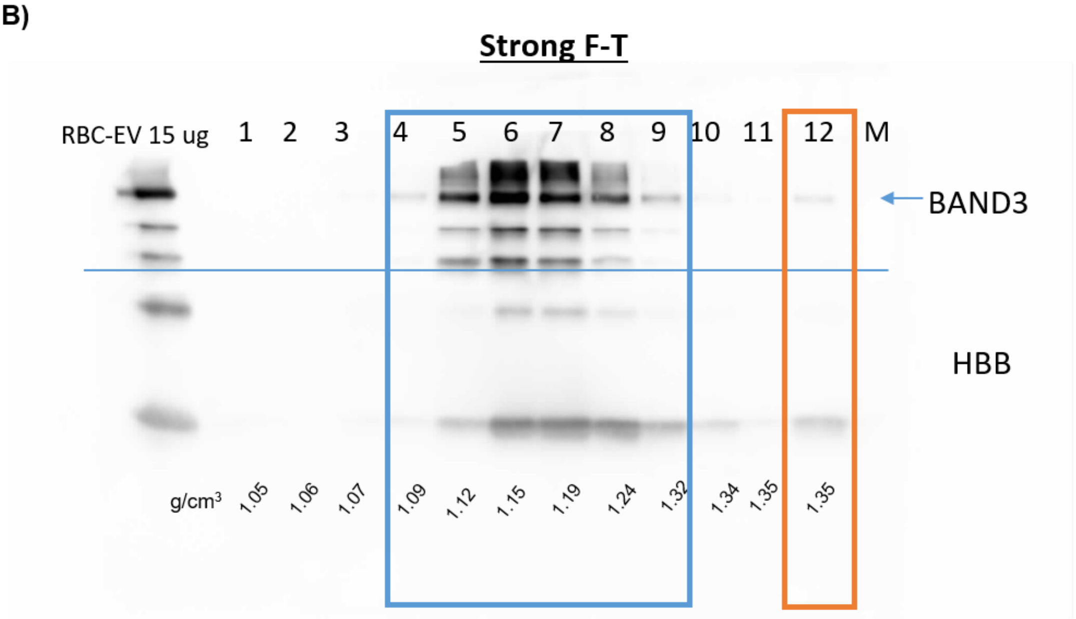


**Figure S6**. Uncropped Western Blot images of gradient distribution of RBC-EVs after Light Freeze and Thaw (Light F-T) (A) and Strong Freeze and Thaw (Strong F-T) (B) treatment.

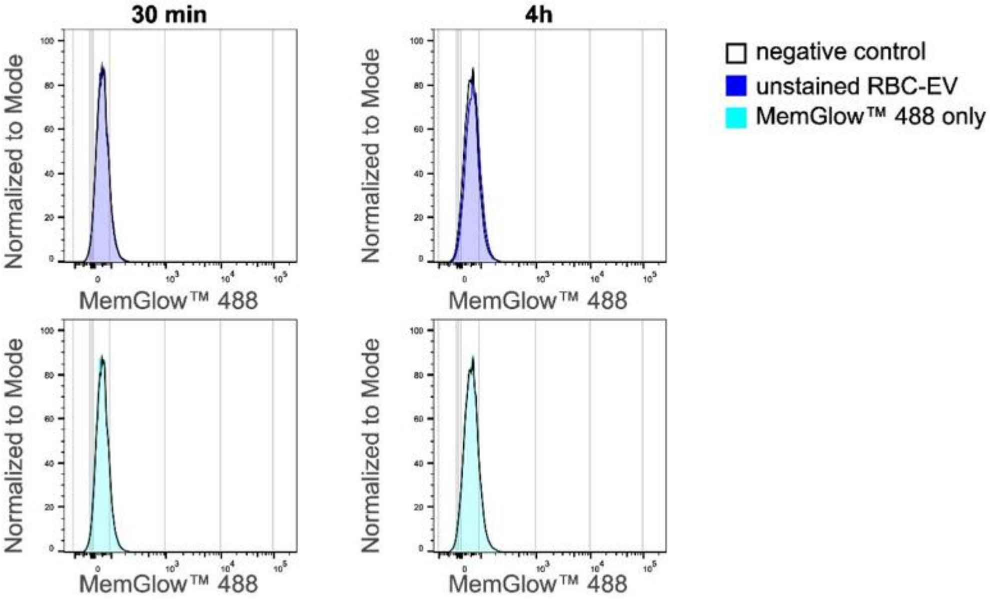


**Figure S7: Flow cytometry analysis of RBC-EV uptake.** Evaluation of unspecific fluorescent signal arising either from possibly increased cell autofluorescence (untreated cells, with histogram), due to EV internalization (unstained RBC-EV, blue hystogram) or to unbound MemGlow™ 488 fluorescent dye (MemGlow only, green histogram) incubated for 30 min or 4 hours with MDA-MB-231 cells.
